# *Shigella* IpaH9.8 limits GBP1-dependent LPS release from intracytosolic bacteria to suppress caspase-4 activation

**DOI:** 10.1101/2022.11.14.516511

**Authors:** Lisa Goers, Kyungsub Kim, Patrick J. Canning, Teagan C. Stedman, Xiangyu Mou, Nadja Heinz Ernst, Jörn Coers, Cammie F. Lesser

**Author notes:** authors contributed equally (Lisa Goers, Kyungsub Kim). **Author Contributions**: LG, KK, XM, CFL designed research; LG, KK, PC, TCS, NHE performed research; LG, KK, PC, TCS, NHE, JC, CFL analyzed data; LG, KK, CFL wrote the paper; KK, CFL, JC obtained funding. **Competing Interest Statement**: No competing interests here. **Classification**: Biological Sciences/Microbiology/Immunology and Inflammation.

## Abstract

Pyroptosis is an inflammatory form of cell death induced upon the recognition of invading microbes. During an infection, pyroptosis is enhanced in interferon-gamma (IFNγ) exposed cells due to the actions of members of the guanylate binding protein (GBP) family. GBPs promote activation of caspase-4 (CASP4) by enhancing its interactions with lipopolysaccharide (LPS), a component of the outer envelope of Gram-negative bacteria. Once activated, CASP4 promotes the formation of non-canonical inflammasomes, signaling platforms that mediate pyroptosis. To establish an infection, intracellular bacterial pathogens, like *Shigella* species, inhibit pyroptosis. The pathogenesis of *Shigella* is dependent on its type III secretion system, which injects ~30 effector proteins into host cells. Upon entry into host cells, *Shigella* are encapsulated by GBP1, followed by GBP2, GBP3, GBP4, and in some cases, CASP4. It has been suggested that the recruitment of CASP4 to bacteria leads to its activation. Here, we demonstrate that two *Shigella* effectors, OspC3 and IpaH9.8, cooperate to inhibit CASP4-mediated pyroptosis. We show that in the absence of OspC3, an inhibitor of CASP4, IpaH9.8 inhibits pyroptosis via its known degradation of GBPs. We find that, while some LPS is present within the host cell cytosol of epithelial cells infected with wild type *Shigella*, in the absence of IpaH9.8, increased amounts are shed in a GBP1-dependent manner. Furthermore, that additional IpaH9.8 targets, likely GBPs, promote CASP4 activation, even in the absence of GBP1. These observations suggest that by promoting LPS release, GBP1 provides CASP4 access to LPS, thus promoting host cell death via pyroptosis.

**Significance Statement:** Non-canonical inflammasomes mediate the death of intestinal epithelial cells in response to invading Gram-negative bacterial pathogens. CASP4 is activated upon binding to LPS. Recent studies have suggested that GBP1 recruits and enables the activation of CASP4 on the surface of intracytosolic bacteria. This has led to the proposal that non-canonical inflammasomes assemble on the bacterial surface due to GBP1-mediated access of CASP4 to the membrane-embedded lipid A component of LPS. Here, we show that by targeting the degradation of GBPs, *Shigella* IpaH9.8 limits the release of LPS from intracellular *Shigella* into the host cell cytosol in a GBP1-dependent manner. Furthermore, we find evidence to suggest that in the absence of GBP1, other members of the GBP family promote CASP4 activation.

## Introduction

Host cell death via pyroptosis is a major arm of the human innate immune response and one of the host’s first lines of defense against invading microbial pathogens. Pyroptosis is an inflammatory lytic form of cell death that results in the loss of a niche for intracellular pathogens and the release of pro-inflammatory cytokines (1). Pyroptosis is triggered by the formation of inflammasomes, multi-protein complexes that assemble upon recognition of cytosolic pathogen-associated molecular patterns (PAMPs), including lipopolysaccharide (LPS) (2–4), the outer component of the cell envelope of Gram-negative bacteria. The assembly of inflammasomes leads to the activation of inflammatory caspases that cleave gasdermin D (GSDMD), a pore-forming protein whose N-terminal domain inserts into and forms pores in the plasma membrane of host cells, eventually leading to their lysis (5–7).

Infection with enteric Gram-negative pathogens primarily triggers pyroptosis via CASP4 activation (8–11). CASP4 is activated upon binding to the hydrophobic membrane-embedded lipid A portion of lipopolysaccharide LPS (4). As an infection progresses and the host responds, intestinal epithelial cells are exposed to cytokines, including interferon-gamma (IFNγ), which induce the expression of hundreds of genes, including guanylate binding proteins (GBPs). GBPs play numerous roles in cell-autonomous immunity, including promoting the recognition of cytosolic LPS by CASP4 (12–15).

Upon entering the cytosol of IFNγ-primed epithelial cells, bacteria are encapsulated by GBPs. GBP1 is the first to bind via direct interactions with LPS (16, 17). Thousands of molecules of GBP1 intercalate into the outer bacterial membrane to form a stable coat, referred to as the GBP1 microcapsule or coatomer (16, 18). Subsequently, GBP2, GBP3, and GBP4 (19–21), and in some cases, CASP4 (10, 16, 17, 22) are recruited to the bacteria. These observations led to the proposal that GBP1 provides CASP4 access to lipid A, thus enabling the assembly of CASP4 platforms on the surface of Gram-negative bacteria (10, 17).

*Shigella* species, the causative agents of bacillary dysentery, are a leading contributor to diarrheal mortality worldwide (23, 24). *Shigella* spp. establish a replicative niche within the cytosol of intestinal epithelial cells and spread from cell to cell using actin-based motility. The virulence of *Shigella*, like many other Gram-negative bacterial pathogens, is dependent on a type III secretion system (T3SS). T3SSs are complex membrane-embedded nanomachines that serve as a conduit for the direct transfer of bacterial proteins, referred to as effectors, into the cytosol of host cells. *Shigella* secrete ~30 unique effectors, including OspC3. OspC3 directly post-translationally modifies and inhibits CASP4 to inhibit pyroptosis (8, 10, 25, 26).

In this study, we demonstrate that *Shigella* IpaH9.8, an effector that targets the ubiquitination and degradation of GBPs (19–21), cooperates with OspC3 to limit *Shigella-*triggered pyroptosis. We demonstrate that while intracellular *Shigella* release LPS into the cytosol of host cells, in the absence of IpaH9.8, these levels are increased 3-5-fold in a GBP1-dependent manner. Furthermore, we find evidence for GBP1-independent pyroptotic cell death of IFNγ-primed epithelial cells in response to infection with *Shigella* that lack OspC3 and IpaH9.8 but not with *Shigella* deficient for only OspC3. These observations demonstrate that GBP1 can be dispensable for CASP4 activation in response to invading bacteria. Furthermore, they suggest that GBP1 promotes LPS release from bacteria into the host cell cytosol, after which GBP2 and/or GBP4, IFNγ-induced targets of IpaH9.8, expressed in intestinal epithelial cells, accelerate pyroptosis due to the activation of non-canonical inflammasomes.

## Results

### Infection with *Shigella* strains that lack OspC3 but not IpaH9.8 triggers pyroptosis of IFNγ-primed epithelial cells

*Shigella* species are intracellular pathogens that establish a replicative niche within the cytosol of colonic epithelial cells, spreading from cell to cell using actin-based motility. To do so, it is essential that intracellular pathogens, like *Shigell*a, inhibit pyroptotic host cell death. We, and others, previously reported that *Shigella* OspC3 suppresses cell death of infected naive untreated epithelial cells via inhibition of CASP4 (8, 10, 25, 27). When we repeated these studies with IFNγ-primed epithelial cells, we observed significantly accelerated rates of host cell death as monitored by uptake of propidium iodide (PI), a phenotype previously reported by others (10, 27).

PI, a membrane-impermeable dye that rapidly enters cells via GSDMD pores, is commonly used as a real-time reporter of pyroptosis. We observed similar rates of PI-uptake using plate reader or automated microscopy-based assays. An advantage of the latter approach was that we could quantify the percentage of pyroptotic cells by analyzing images of cells exposed to both PI and Hoechst, a membrane-permeable dye taken up by all cells, dead or alive. In each case, after providing time for the bacteria to invade host cells, gentamicin was added to kill any that remained extracellular.

Under all conditions tested, regardless of the presence of AfaI, an adhesin that promotes host cell attachment and synchronizes host cell invasion (28), infection with Δ*ospC3 Shigella* triggered significant levels of cell death of IFNγ-primed HeLa (Fig. 1A-B, Fig. S1A) and HCT8 cells (Fig. 1C, Fig S1C). In each case, cell death depended on GSDMD (Fig. 1B-C, Fig. S1D), establishing that PI-uptake in these assays reflected activation of pyroptosis. We conducted all the following experiments with AfaI-expressing *Shigella*.

**Figure 1:**
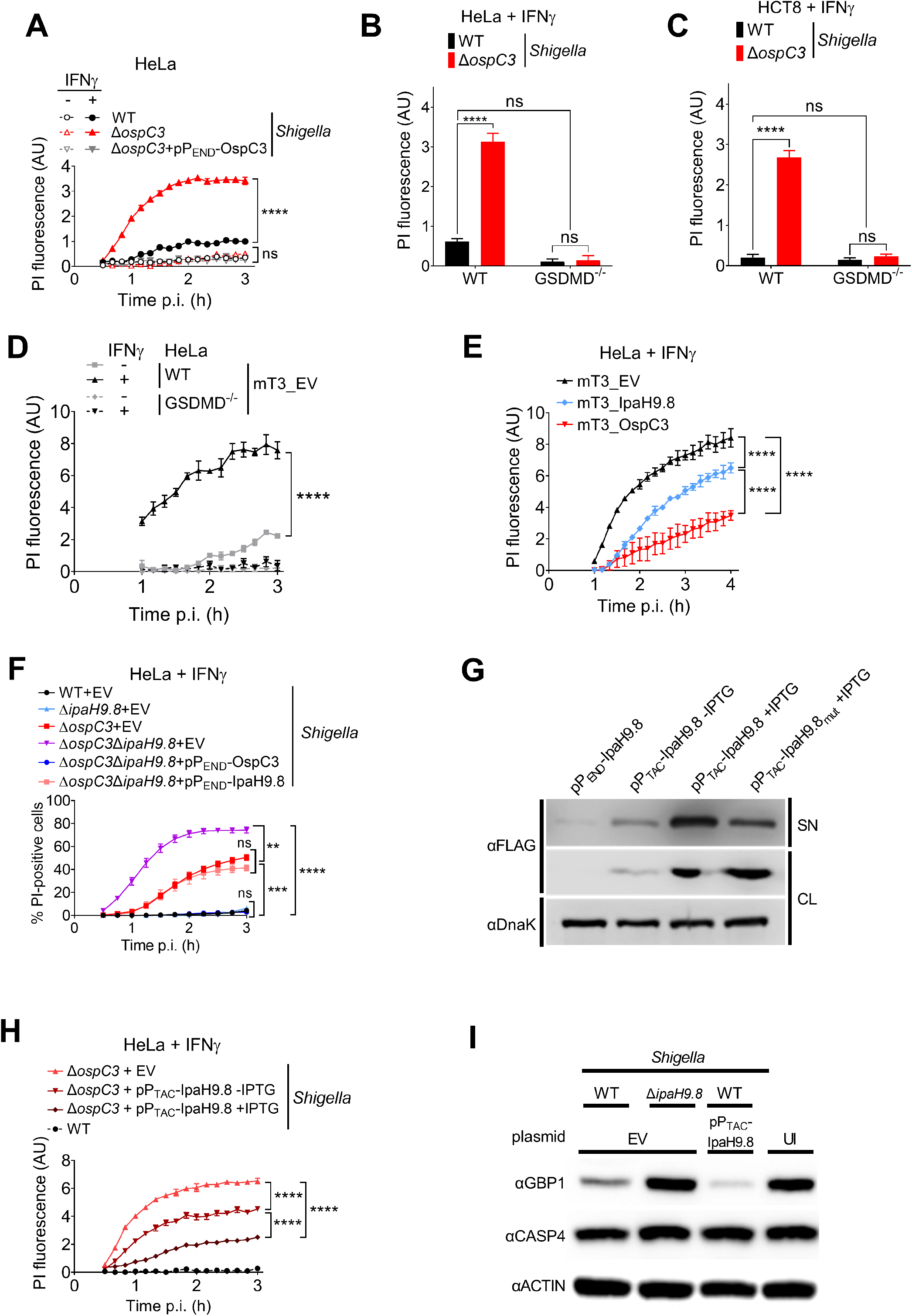
*Shigella* OspC3 and IpaH9.8 cooperate to suppress bacterial-triggered pyroptosis of IFNγ-primed epithelial cells. (A-F, H, I) WT or GSDMD^−/−^ HeLa or HCT8 cells unprimed or primed overnight with 10 ng/ml IFNγ were infected with the indicated strains at an MOI of 100 (A), 10 (B-F, H), or 5 (I). Cells were infected with *Shigella* strains that carry pBAD33-AfaI and designated plasmids (B-C, H-I). For *Shigella*, at thirty minutes post-infection (p.i.) (A-C, F, H, I) and, for mT3, at 1-hr p.i. (D, E), PI (A-E, H) or PI and Hoechst (F) were added to the medium, and cell death was measured by monitoring PI-uptake using a plate reader (A-E, H) or PI and Hoechst uptake using an automated fluorescence microscope (F). A time course of cell death is shown in (A, D-F) and a 3-hour endpoint in (B). (G) Immunoblots of secretion assays of IpaH9.8-FLAG expressed from designated plasmids by Δ*ospC3 Shigella*. SN (supernatant) and CL (cell lysate) fractions were probed with anti-FLAG to detect secreted and expressed effectors, and DnaK, a cytosolic *Shigella* protein, as a lysis and loading control. (I) Immunoblots of lysates of IFNγ-primed HeLa cells infected with indicated *Shigella* strains probed with indicated antibodies. EV= empty vector, UI = uninfected, and AU= arbitrary units. Values shown are the mean +/− SEM of three experimental replicates. Data were analyzed using two-way ANOVA with Tukey’s post hoc test. *P < 0.05, **P < 0.01, ***P < 0.001, ****P < 0.0001, ns = non-significant.

GBPs are among some of the most highly induced genes upon exposure of epithelial cells to IFNγ and have been shown to promote activation of CASP4 inflammasomes. Multiple of the members of this family, including human GBP1, GBP2, GBP4, and GBP6, are targeted for degradation by *Shigella* IpaH9.8 (19). However, like others (20, 27), we observed no evidence of cell death of IFNγ-primed HeLa or HCT8 cells infected with Δ*ipaH9.8 Shigella* (Fig. S1E-F) under conditions whereby, as expected, we detected IFNγ-dependent GBP1 expression (Fig. S1G).

#### OspC3 and IpaH9.8 suppress mT3Ec-triggered cell death of IFNγ-primed epithelial cells

Type III secreted effectors often act in an additive or functionally redundant manner to target host cell processes such that phenotypes associated with their absence from bacteria are not detected using traditional top-down approaches with single deletion strains. Reasoning that IpaH9.8 acts in concert with other effectors to suppress cell death, we investigated whether we could identify a role for IpaH9.8 in regulating cell death using a bottom-up platform.

mT3.1 *E. coli*, referred to herein as mT3Ec, is a variant of non-pathogenic DH10β *E. coli* that expresses the *Shigella Mxi, Spa*, and *Inv* operons. These operons encode the structural components needed to form the *Shigella* type III secretion apparatus (T3SA) plus four embedded effectors (29). mT3Ec, like WT *Shigella*, invades and efficiently escapes into the cytosol of epithelial cells (29). In contrast to WT *Shigella*, mT3Ec invasion into epithelial cells triggers host cell death via pyroptosis (8).mT3Ec can secrete the full complement of *Shigella* effectors (8). In a prior study with unprimed epithelial cells, we found that adding OspC3, but not IpaH9.8,inhibits mT3Ec-triggered cell death (8).

Given that IFNγ-priming resulted in enhanced Δ*ospC3 Shigella-*triggered cell death, we reasoned that IFNγ-primed cells would exhibit a more robust cell death response when infected with mT3Ec. This was the case, as we observed enhanced mT3Ec-triggered cell death of IFNγ-primed as compared to unprimed HeLa and HCT8 epithelial cells, which was dependent on GSDMD (Fig.1D, Fig. S1H). This demonstrates that infection with mT3Ec, like Δ*ospC3 Shigella*, triggers pyroptosis of IFNγ-primed epithelial cells.

We next compared the PI-uptake of IFNγ-primed cells infected with mT3Ec that contain an empty plasmid (mT3Ec_EV) to variants that express OspC3 (mT3Ec_OspC3) or IpaH9.8 (mT3Ec_IpaH9.8). The addition of plasmids that express each effector under the control of an isopropyl β-D-1-thiogalactopyranoside (IPTG)-inducible P_TAC_ promoter (pP_TAC_-OspC3 or pP_TAC_-IpaH9.8) led to significantly diminished mT3Ec-triggered cell death, OspC3 to a greater extent than IpaH9.8 (Fig. 1E). These observations suggested that IpaH9.8, like OspC3, plays a role in inhibiting *Shigella-*triggered pyroptosis.

#### OspC3 and IpaH9.8 cooperate to inhibit *Shigella*-triggered cell death of IFNγ-primed epithelial cells

As we observed no evidence for IpaH9.8 in suppressing pyroptosis when absent from otherwise WT *Shigella*, we reasoned that IpaH9.8-mediated GBP degradation acts upstream of OspC3 to limit CASP4 activation. Thus, we compared the fate of IFNγ-primed epithelial cells infected with WT, Δ*ospC3*, or Δ*ospC3*Δ*ipaH9.8 Shigella*. HeLa cells infected with Δ*ospC3*Δ*ipaH9.8* were ~1.5 fold more likely than those infected with Δ*ospC3 Shigella* to trigger PI-uptake (Fig. 1F), thus exhibiting evidence of enhanced cell death via pyroptosis. We found that the introduction of a plasmid that encodes IpaH9.8 under the control of its endogenous promoter (P_END_) reduced Δ*ospC3*Δ*ipaH9.8 Shigella*-triggered cell death to levels equivalent to Δ*ospC3 Shigella*. IFNγ-primed HCT8 cells behaved similarly when infected with the same set of strains (Fig. S1I). Δ*ospC3*Δ*ipaH9.8 Shigella*-triggered cell death was also completely GSDMD-dependent (Fig. S1J).

IpaH9.8 is a “second-wave” *Shigella* effector whose expression is regulated by MxiE; a transcription factor only activated post-contact with host cells (30). To gain additional evidence that IpaH9.8 limits *Shigella-*triggered cell death, we investigated whether we could suppress Δ*ospC3 Shigella*-triggered cell death by increasing the amount and accelerating the timing of translocation of IpaH9.8 into host cells. Thus, we infected IFNγ-primed epithelial cells with Δ*ospC3 Shigella* that carry pP_TAC-_IpaH9.8. The P_TAC_ promoter is leaky but inducible in *Shigella*, such that, even in the absence of IPTG, Δ*ospC3*/P_TAC-_IpaH9.8 *Shigella* secrete more IpaH9.8 than Δ*ospC3*/ P_END-_IpaH9.8, which express IpaH9.8 under the control of its endogenous promoter (Fig. 1G). We observed decreased levels of Δ*ospC3 Shigella*-triggered cell death as secreted levels of IpaH9.8 were increased (Fig. 1H).

In addition to GBPs, IpaH9.8 has also been reported to promote the ubiquitination of NEMO/IKKγ (31) and U2AF (32). To assess whether IpaH9.8 was suppressing cell death via its targeting of GBPs, we next tested whether secretion of IpaH9.8_Y86A/Q88A (IpaH9.8mut) (Fig. 1G), a variant impaired in GBP binding (33), suppressed cell death. This was not the case (Fig. S1K). To confirm that increased IpaH9.8 secretion led to decreased cell death due to increased degradation of GBPs, we compared levels of GBP1 within lysates of cells infected with WT, Δ*ipaH9.8*, and WT/pP_TAC_ -IpaH9.8 *Shigella*. As expected, we observed decreased levels of GBP1 within lysates of cells infected with WT compared to Δ*ipaH9.8 Shigella*, which were further reduced in cells infected with WT/P_TAC_-IpaH9.8 *Shigella* (Fig. 1I). We conducted these studies with cells infected with *Shigella* that encode OspC3 to ensure that any observed differences in GBP1 levels were not reflective of protein loss due to cell lysis. Together these data establish a role for IpaH9.8-mediated GBP degradation in inhibiting *Shigella-*triggered pyroptosis.

#### OspC3 and IpaH9.8 cooperate to promote the growth of intracytosolic *Shigella*

We next expanded our automated microscopy-based cell death assay to image the real-time replication of bacteria within infected epithelial cells. To differentiate between intra- and extracellular bacteria, we infected with *Shigella*, which express superfolder green fluorescent protein (sfGFP) under the control of the arabinose-inducible *araBAD* promoter (P_BAD_). Following treatment with gentamicin to kill extracellular bacteria, we introduced arabinose to induce sfGFP expression (Fig. S2A). Under these conditions, sfGFP-expressing bacteria represent live replicating intracellular bacteria.

Using this assay, we compared the fate of WT, Δ*ospC3*, and complemented Δ*ospC3*/P_END_-OspC3 *Shigella* within untreated and IFNγ-primed epithelial cells. We detected evidence of sfGFP-expressing intracellular bacteria starting at ~60 min post the addition of arabinose (Fig. S2C). We found similar rates of increasing GFP signal/infected cell within untreated epithelial cells infected with WT, Δ*ospC3*, and Δ*ospC3*/P_END_-OspC3 (Fig. 2A), suggesting, as previously shown, that OspC3 is not involved in host cell invasion (8). In contrast, we observed decreased evidence of replication of Δ*ospC3* but not WT or Δ*ospC3*/P_END_-OspC3 *Shigella* within infected IFNγ-primed cells (Fig. 2A). Only infection with Δ*ospC3 Shigella* triggered cell death, as assessed by PI uptake (Fig 2B, Fig. S2B, C).

**Figure 2:**
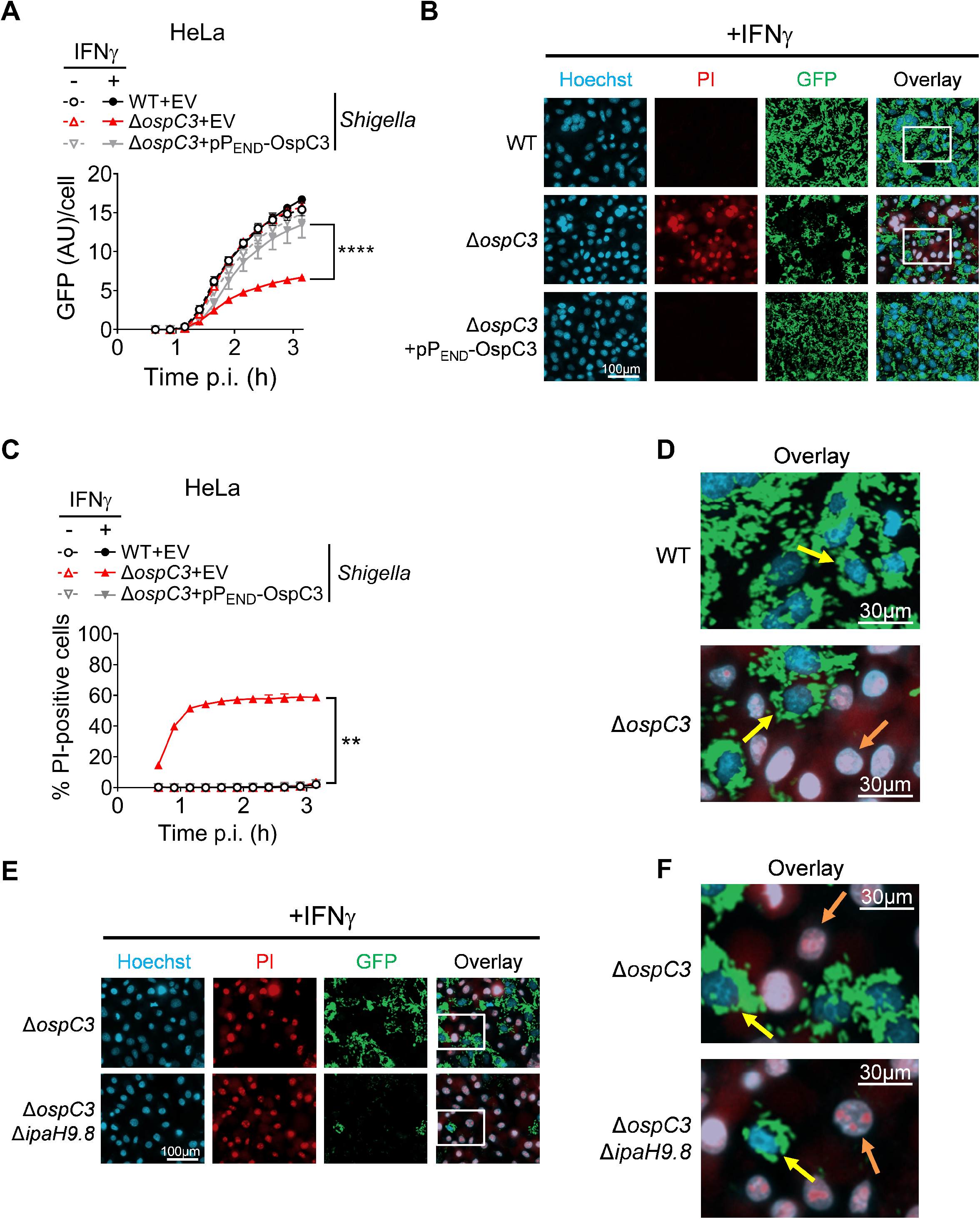
*Shigella* OspC3 and IpaH9.8 cooperate to promote the growth of intracytosolic *Shigella*. (A-F) WT HeLa cells unprimed or primed overnight with 10 ng/ml IFNγ were infected at an MOI of 3 with designated strains, each of which carries pBAD33-sfGFP, pBR322-AfaI, and pPEND-OspC3 or an empty vector (EV). Thirty minutes post-infection (p.i.), PI, Hoechst, and arabinose were added to the medium, and cells were imaged using an automated fluorescent microscope. Time course of total GFP divided by the number of Hoechst-positive (Hoechst^+^) nuclei in (A) and percentage of PI^+^/Hoechst^+^ cells in (B). Values shown are the mean +/− SEM of three experimental replicates. Representative images of cells infected with the designated strains at 3-hr p.i. (C, E) with enlarged inserts are shown in (D, F), PI (red), Hoechst (blue), and sfGFP (green). Yellow arrows point to live PI^−^GFP^+^ cells, while orange arrows point to dead PI^+^GFP^−^cells. Three biological replicates were performed, and representative data are shown. Statistical significance was assessed by a two-way ANOVA with Tukey’s post hoc test. *P < 0.05, **P < 0.01, ***P < 0.001, ns = non-significant.

Representative images of IFNγ-primed cells infected with each strain acquired at 3 hours post-infection revealed the existence of two distinct populations of Δ*ospC3 Shigella* infected cells: PI^+^ cells with no evidence of sfGFP^+^ *Shigella* and PI^−^ cells populated with sfGFP bacteria. The number of sfGFP^+^ Δ*ospC3 Shigella* observed within live (PI^−^) cells was similar to that observed within cells infected with WT *Shigella* (Fig. 2B, D). While we were unable to detect bacteria within the pyroptotic PI^+^ cells, bacteria likely invaded these cells, as, under these experimental conditions (MOI 10), we found sfGFP^+^ WT and Δ*ospC3 Shigella* within >95% of unprimed infected cells (Fig. S2D).

We next compared the fate of Δ*ospC3* and Δ*ospC3*Δ*ipaH9.8 Shigella* within infected cells. As before (Fig. 1F), at 3 hours post-infection, we observed increased cell death of IFNγ-primed cells infected with Δ*ospC3*Δ*ipaH9.8* vs. Δ*ospC3 Shigella* (Fig. 2E-F). Again, we found that some infected cells evaded host cell death. In this case, we observed fewer PI^−^ GFP^+^ cells infected with Δ*ospC3*Δ*ipaH9.8* compared to Δ*ospC3 Shigella* (Fig. S2E).

These data are consistent with prior studies, which found that replication of cytosolic *Shigella* and *Salmonella* is impaired within pyroptotic epithelial cells (10, 17).Although, whether bacterial killing reflects an active host cell process versus an experimental artifact of gentamicin uptake via GSDMD pores remains to be determined. Nevertheless, these studies demonstrate that *Shigella* inhibits host cell death to establish a productive replicative niche within IFNγ-primed epithelial cells via the concerted efforts of OspC3 and IpaH9.8.

#### Intracytosolic *Shigella* inhibit pyroptosis in an IpaH9.8-dependent manner

Our finding that a subset of Δ*ospC3 Shigella-*infected epithelial cells evaded cell death was unexpected, as it was previously reported that all host cells infected with Δ*ospC3 Shigella* rapidly undergo pyroptosis (10). A significant difference between our experimental design and the earlier study was the multiplicity of infection (MOI) studied. Thus, we compared the fate of primed epithelial cells infected with Δ*ospC3 Shigella* at an MOI of 3 versus 10. Strikingly, with the lower MOI, we observed a higher percentage of PI^+^ pyroptosing cells (Fig. 3A, C) and decreased bacterial GFP fluorescence (Fig. 3B, C).

**Figure 3:**
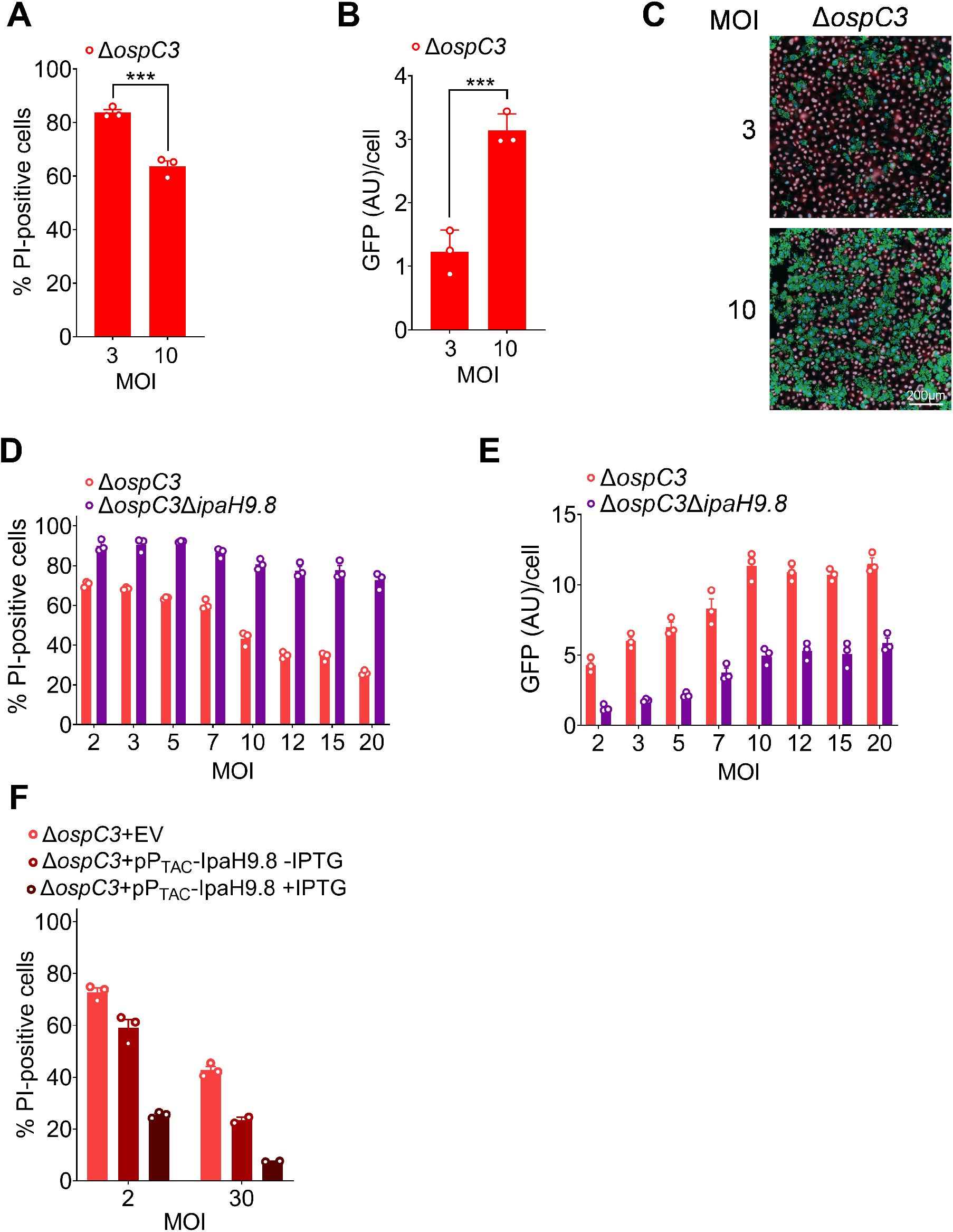
*Shigella* inhibits pyroptosis in an IpaH9.8-dependent manner. (A-F) WT HeLa cells primed with 10 ng/ml IFNγ overnight were infected with designated strains at the specified MOIs. Strains carry either pBAD33-sfGFP and pNG162-AfaI (A-E) or pBAD33-AfaI and the designated plasmids (F). Thirty minutes post-infection, PI, Hoechst, and arabinose were added to the medium, after which cells were imaged using an automated fluorescent microscope. Quantification of % PI-positive cells (PI^+^/Hoechst^+^ cells) at 1.5 h p.i. (A, C, F) and GFP fluorescence/Hoechst^+^ cell at 3 h p.i. (B, E). Representative images of cells infected with Δ*ospC3 Shigella* at designated MOIs at 3 h p.i. (C). Values shown are the mean +/− SEM of three experimental replicates. Three biological replicates were performed, and representative data are shown. For (A) and (B), single time points of a time course are shown, and statistical significance was assessed by an ordinary one-way ANOVA. *P < 0.05, **P < 0.01, ***P < 0.001, ****P < 0.0001, ns = non-significant.

We expanded these studies to include both Δ*ospC3* and Δ*ospC3*Δ*ipaH9.8 Shigella*, testing MOIs from 2 to 20. We observed decreasing evidence of pyroptosing cells with increasing MOIs (Fig. 3D, Fig. S3A, B). This phenomenon was more striking when infecting with Δ*ospC3* than Δ*ospC3*Δ*ipaH9.8 Shigella*, i.e., 65 to 20% versus 90 to 80% cell death (Fig. 3D, Fig. S3A). We also observed increased evidence of intracytosolic bacterial replication as the MOI was raised (Fig. S3C-D), with a direct correlation between detectable GFP fluorescence and MOI (Fig. 3E).

These observations suggested that a significant driving factor in determining whether infected host cells undergo rapid cell death upon *Shigella* invasion is the presence of IpaH9.8. To directly test this hypothesis, we infected the cells with Δ*ospC3*/P_TAC_-IpaH9.8 that secrete excess IpaH9.8 (Fig. 1G). Regardless of MOI, we found that Δ*ospC3 Shigella-*triggered host cell death decreased as levels of IpaH9.8 were increased (Fig. 3F). Remarkably, at an MOI of 30, IPTG-induced expression of IpaH9.8 reduced Δ*ospC3 Shigella* triggered cell death to almost undetectable levels. These data suggest significant roles for GBP1, GBP2, GBP4, and/or GBP6, the targets of IpaH9.8, in promoting rapid host cell death via CASP4-mediated pyroptosis.

### GBP1 enhances LPS release from intracytosolic *Shigella*

We next investigated the molecular mechanism by which IpaH9.8 limits *Shigella-*triggered pyroptosis. Upon invasion into host cells, *Shigella* are encapsulated by GBP1, followed by GBP2, GBP3, and GBP4. GBP1 polymers bind to the exposed outer O-antigen polysaccharide portion of LPS on the surface of the bacteria and then depolymerize into a bacteria-engulfing microcapsule (16, 20), which acts as an LPS surfactant to destabilize the rigidity of the outer bacterial membrane (16). In the case of *Shigella*, this binding not only impairs IcsA localization and actin tail formation but also increases the *in vitro* sensitivity of *Shigella* to polymixin B, an antimicrobial peptide that targets the outer bacterial envelope (16). As GBP1 promotes destabilization of the outer membrane, we hypothesized that its binding enhances the release of bacterial LPS into the cytosol of host cells.

To address this possibility, we developed an assay to monitor cytosolic LPS (cLPS) levels, adapted from one previously designed to monitor the uptake of extracellular bacterial outer membrane vesicles (OMVs) into the host cell cytosol (34). Two hours post-infection, we exposed *Shigella*-infected epithelial cells to digitonin under conditions that selectively lyse the host cell plasma membrane (35).The cytosol and residual fractions were separated via a low-speed spin, after which we filtered the cytosolic fractions to remove any remaining bacteria. *Shigella* in the residual and filtered supernatant fractions were enumerated, and LPS levels in the filtered cytosolic fractions were quantified using a Limulus amoebocyte lysate (LAL) assay. We assessed fractionation efficiency and GBP1 levels by immunoblotting for GBP1, caveolin, a plasma membrane protein, GroEL, a cytosolic *Shigella* protein, and actin.

Given that the absence of IpaH9.8 is associated with impaired *Shigella* cell-to-cell spread within IFNγ-primed epithelial cells due to delayed actin tail formation (20, 21), we conducted these studies with variants of *Shigella* that lack IcsA, i.e., Δ*icsA* and Δ*icsA*Δ*ipaH9.8 Shigella*. Infection with neither of these OspC3-containing strains triggered pyroptosis of IFNγ-primed epithelial cells (Fig. S4A). We also infected cells with non-invasive virulence-plasmid-minus BS103 *Shigella* (20, 21) to assess whether host cell invasion is needed to detect cLPS.

We first compared cLPS levels in lysates of IFNγ-primed and unprimed epithelial cells infected with Δ*icsA*, Δ*icsA*Δ*ipaH9.8*, or BS103 *Shigella* (Fig. 4A). We observed similar levels of cLPS within lysates of unprimed cells infected with Δ*icsA* and Δ*icsA*Δ*ipaH9.8 Shigella*. Strikingly, we detected ~5-fold higher levels of cLPS within lysates of IFNγ-primed cells infected with Δ*icsA*Δ*ipaH9.8* compared to Δ*icsA Shigella*, while the levels of cLPS present in the Δ*icsA Shigella* infected cells were equivalent to those found in unprimed cells. As expected, we found diminished levels of GBP1 in the lysates of primed cells infected with Δ*icsA* as compared to Δ*icsA*Δ*ipaH9.8 Shigella* (Fig. 4B). Equivalent numbers of Δ*icsA* and Δ*icsA*Δ*ipaH9.8 Shigella* were present within the residual fractions (Fig. 4C). No GBP1 was detected in unprimed cells (Fig. 4B) and no bacteria found in the filtered cytosolic fractions (Fig. 4C). Together these data demonstrate that intracellular *Shigella* shed LPS into the host cell cytosol of unprimed cells, which is substantially enhanced in IFNγ-primed cells due to the action of one or more IFNγ-induced targets of IpaH9.8 that are expressed in intestinal epithelial (GBP1, GBP2, and/or GBP4).

**Figure 4:**
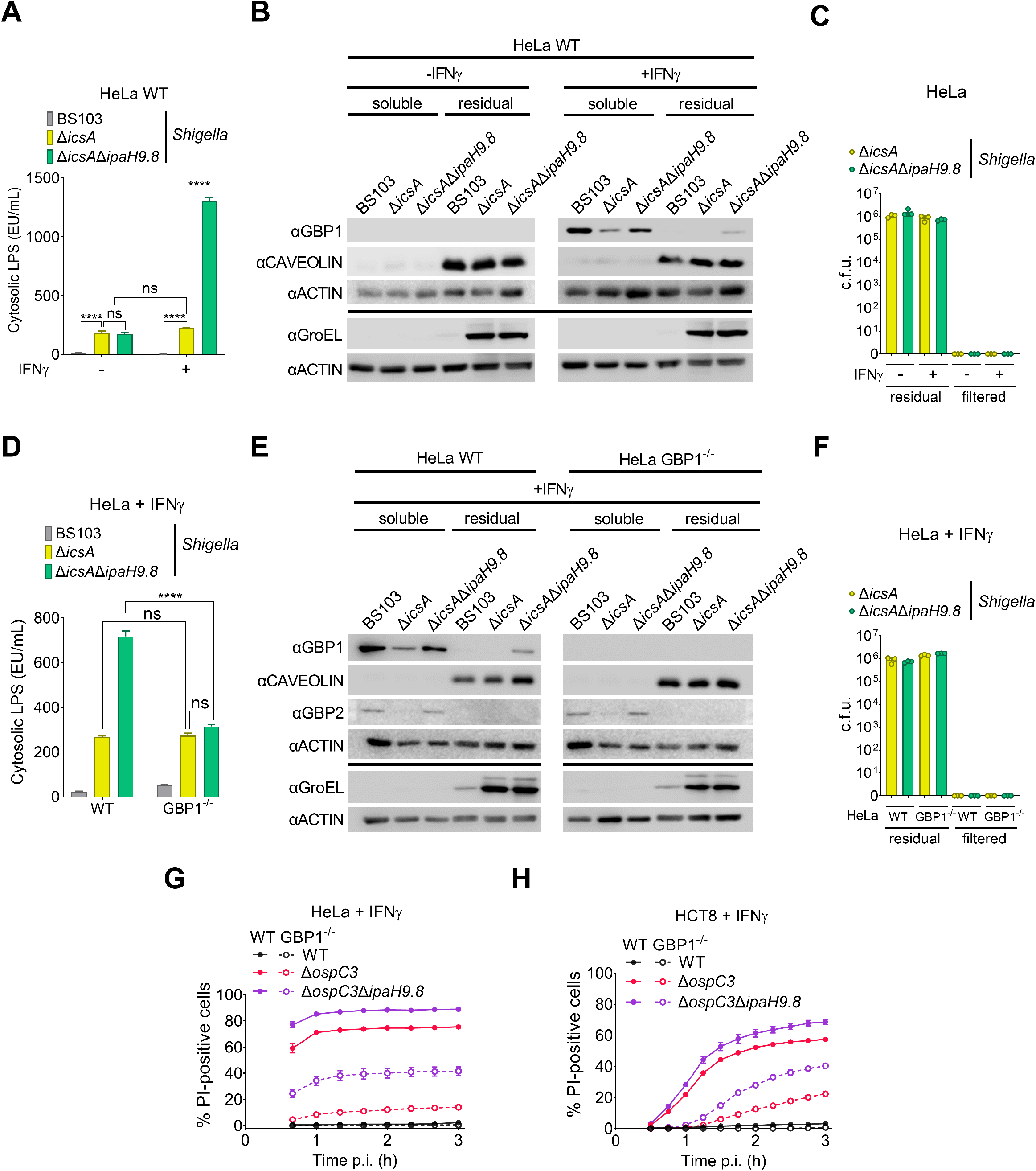
GBP1 enhances LPS release from intracellular *Shigella* but is not essential for CASP4-mediated pyroptosis of infected cells. (A-H) WT or GBP1^−/−^ HeLa or HCT8 cells unprimed or primed overnight with 10 ng/ml IFNγ were infected with the indicated *Shigella* strains that each carry pNG162-AfaI at an MOI of 3. (A-F) Two hours post-infection, cells were lysed with 0.005% digitonin and separated into cytosolic and residual fractions. LPS levels in filtered cytosolic extracts were quantified using a LAL assay (A, D), fractionation efficiency and GBP1 levels were assessed by immunoblotting fractions with designated antibodies (B, E), and numbers of bacteria within the residual and filtered supernatant fractions were enumerated (C, F). (G-H) Thirty minutes post-infection, gentamicin, PI, and Hoechst were added to the medium, and cells were imaged using an automated fluorescent microscope. Time courses of cell death PI^+^/Hoechst ^+^ cells are shown. Values shown are the mean +/− SEM of three experimental replicates. Where indicated, statistical significance was assessed using two-way ANOVA with Tukey’s post hoc test. *P < 0.05, **P < 0.01, ***P < 0.001, ****P < 0.0001, ns = non-significant.

To investigate a direct role for GBP1 in mediating LPS release, we generated two independent GBP1^−/−^ HeLa cell lines (GBP1^−/−^ 1-9, GBP1^−/−^ 1-10) using CRISPR/Cas9 editing. After confirming that each line still expressed its closest homolog, GBP2 (Fig. S4B), we compared the levels of cLPS present within the parental and modified IFNγ-primed cell lines infected with BS103 and Δ*icsA*Δ*ipaH9.8 Shigella*. For each of the two GBP1^−/−^ cell lines, we observed an ~3-fold decrease in detectable cLPS (Fig. S4C). We extended these studies and compared cLPS levels within lysates of IFNγ-primed WT and GBP1^−/−^ cells infected with BS103, Δ*icsA*, and Δ*icsA*Δ*ipaH9.8 Shigella*. We again observed substantially lower levels of cLPS within GBP1^−/−^ as compared to WT cells infected with Δ*icsA*Δ*ipaH9.8 Shigella* (Fig. 4D). As expected, we found diminished levels of GBP1 and GBP2 in the lysates of primed cells infected with Δ*icsA* as compared to Δ*icsA*Δ*ipaH9.8 Shigella* and none in GBP1^−/−^ cells (Fig. 4E). Equivalent numbers of Δ*icsA* and Δ*icsA*Δ*ipaH9.8 Shigella* were present within the residual fractions (Fig. 4F). Strikingly, we detected equivalent levels of cLPS in lysates of IFNγ-primed WT cells infected with Δ*icsA Shigella* and GBP1^−/−^ cells infected with Δ*icsA* and Δ*icsA*Δ*ipaH9.8 Shigella*. Together, these data suggest that there is a baseline release of LPS from intracytosolic *Shigella*, which is enhanced in a GBP1-dependent manner.

### GBP1 promotes but is not essential for the activation of pyroptosis in response to invading bacteria

Given that GBP1 promotes LPS release from *Shigella* and LPS promotes CASP4 activation, we next compared the fate of IFNγ-primed WT and GBP1^−/−^ HeLa cells when infected with WT, Δ*ospC3*, and Δ*ospC3*Δ*ipaH9.8 Shigella*. Consistent with prior studies with IFNγ-primed HeLa cells (10), we observed almost complete suppression of Δ*ospC3 Shigella*-triggered cell death in the absence of GBP1 (Fig. 4G). However, in contrast, we observed only ~50% suppression of Δ*ospC3*Δ*ipaH9.8 Shigella*-triggered cell death, which was suppressed to almost undetectable levels in the presence of disulfiram, a small molecule inhibitor of GSDMD (36) (Fig. S4D).

We also compared the fate of WT and GBP1^−/−^ HCT8 cells (Fig. S4E) infected with WT, Δ*ospC3*, and Δ*ospC3*Δ*ipaH9.8 Shigella*. In this case, we again observed significant suppression of Δ*ospC3* and moderate suppression of Δ*ospC3*Δ*ipaH9.8 Shigella-*triggered cell death in the absence of GBP1 (Fig. 4H), which were further suppressed in the presence of disulfiram (Fig. S4F). The major difference between GBP1^−/−^ cells infected with Δ*ospC3* versus Δ*ospC3*Δ*ipaH9.8 Shigella* is that those infected with bacteria that inject Ipa9.8 (Δ*ospC3* Shigella) will have significantly lower levels of GBP2, GBP4, and GBP6. Together, these studies demonstrate that GBP1 is not essential for the activation of non-canonical CASP4 inflammasomes and suggest that other IFNγ-induced targets of IpaH9.8 that are expressed in intestinal epithelial cells, e.g., GBP2 and/or GBP4, promote CASP4 activation, even in the absence of GBP1.

## Discussion

To establish a replicative niche within the cytosol of human intestinal epithelial cells, it is essential for invading pathogens to evade host cell death via pyroptosis, which in the case of Gram-negative bacterial pathogens, is induced upon CASP4 sensing of LPS, a response that is enhanced in IFNγ-primed cells. Upon entry into the cytosol of IFNγ-primed cells, bacteria are rapidly engulfed in a GBP1 microcapsule (16). GBP2, GBP3, and GBP4 are recruited, followed by CASP4, suggesting that GBPs promote the assembly of CASP4 inflammasomes on the surface of bacteria (10, 17).

*Shigella* IpaH9.8 targets multiple human GBPs for degradation, e.g., GBP1, GBP2, GBP4, and GBP6. Previous studies have shown that in the absence of IpaH9.8, CASP4 is recruited to a subset of intracytosolic *Shigella* (10, 16). In this study, using a variety of complementary approaches, we have established that IpaH9.8 acts upstream of OspC3 to suppress *Shigella-* triggered cell death. Using a quantitative LAL assay, we found that *Shigella* releases LPS into the cytosol of unprimed host cells. In the absence of IpaH9.8, within IFNγ-primed cells, these levels are increased 3-5-fold in a GBP1-dependent manner. Consistent with this hypothesis, Zhu et al. found that GBP1 enhances the release of labeled LPS from intracellular *S*. Typhimurium using a fluorescent microscopy-based assay (18). These observations suggest that GBP1 mediates LPS release from intracellular bacteria into the host cell cytosol; whether the liberated LPS is in clusters or within OMVs will be the subject of future studies.

It has been proposed that GBP1 is essential for the activation of CASP4 inflammasomes. Yet, there is extensive evidence of CASP4 activation in response to the invasion of Gram-negative bacteria into unprimed (3, 8, 11) and GBP-deficient IFNγ-primed epithelial cells (27). Based on our observation that *Shigella* shed LPS into the cytosol of even in unprimed epithelial cells, we propose that spontaneously shed LPS, which increases as *Shigella* replicate within the host cell cytosol, directly binds to and activates CASP4, even in the absence of GBPs, likely accounting for the delayed onset of pyroptosis observed in response to invading *Shigella* in unprimed WT and IFNγ-primed epithelial GBP deficient epithelial cells.

As previously reported, we also found that infection with Δ*ospC3 Shigella* triggers substantially lower levels of cell death of IFNγ-primed GBP1^−/−^ HeLa and HCT8 cells as compared to their WT parental cell lines. However, the absence of GBP1 only results in moderate suppression of Δ*ospC3*Δ*ipaH9.8 Shigella* triggered cell death. As Δ*ospC3*Δ*ipaH9.8 Shigella* does not translocate IpaH9.8 into host cells, IFNγ-primed epithelial cells infected with this strain, as compared to those infected with Δ*ospC3 Shigella*, have detectable GBP2 and GBP4 (19). Thus, we propose that in the absence of GBP1, the relatively low levels of LPS that are spontaneously shed into the host cell cytosol interact with GBP2 and/or GBP4, perhaps in concert with GBP3 which was previously demonstrated to promote CASP4 activation (17), to promote pyroptosis in a GBP1-independent manner. Consistent with this hypothesis, Dickinson et al. recently found that GBP2, like GBP1, upon direct binding to LPS, polymerizes with LPS into large complexes (37), perhaps forming cytosolic platforms for the assembly of non-canonical CASP4 inflammasomes.

Interestingly, we found that IpaH9.8-mediated GBP degradation plays an increasing role in promoting cell death as the initial numbers of intracytosolic *Shigella* increase. It is difficult to determine what is a physiologically relevant MOI, particularly after the establishment of an infection. Presumably, higher titers of *Shigella* are likely located within the cytosol of intestinal epithelial cells at later points in an infection when the cells are exposed to IFNγ. Regardless, it is important to note that changes in MOI can lead to vastly different conclusions about the roles of virulence proteins.

In summary, here we present complementary experimental approaches that *Shigella* IpaH9.8 modulates cell death via its targeted degradation of GBPs. We show that IpaH9.8 limits LPS release from intracellular *Shigella* in a GBP1-dependent manner and that GBP1 is not essential for at least the early onset of *Shigella-*triggered cell death. These observations suggest that the recruitment of CASP4 to the surface of bacteria via GBPs is not essential for its activation but rather that GBP1 promotes the release of LPS into the host cell cytosol, whereby interactions with additional GBPs promote CASP4 activation. Given that GBP1 promotes pyroptosis in response to transfected LPS and cytosolic OMVs, it is likely that it also acts in the recognition of LPS released from intracellular bacteria (12, 15), a topic for future investigation.

## Materials and Methods

### Bacterial strains and plasmids

All strains used are listed in Table S1, and plasmids in Table S2.

#### Generation of IpaH9.8 expression plasmids

pP_END_-IpaH9.8 and pP_TAC_-IpaH9.8mut were each generated using Gateway™ cloning. For pP_END_-IpaH9.8_mut_, a pENTR221 entry clone was developed using a PCR-amplified fragment of the *Shigella* virulence plasmid that contains IpaH9.8 plus 295 upstream nucleotides generated using oligos END98-5 and END98-3 (Table S3). For P_TAC_-IpaH9.8_mut_, a pENTR221 entry clone was developed using a synthetic DNA fragment containing the ORF encoding IpaH9.8_Y86A/Q88A plus an upstream consensus RBS (ribosome binding sequence) flanked by *attB* sites (Twist Bioscience). Once sequence-verified, the introduced regions were introduced into pCMD136-ccdB-FLAG and pDSW206-ccdB-FLAG, respectively.

#### Mammalian cell culture

HeLa and HCT8 cells obtained from the American Type Culture Collection were cultured in a 5% CO_2_ incubator at 37°C. HeLa cells were maintained in high glucose DMEM (Thermo Fisher Scientific #11965) and HCT8 cells in GlutaMAX-supplemented RPMI 1600 (Thermo Fisher Scientific #61870127). In both cases, media were supplemented with 10% heat-inactivated fetal bovine serum (FBS, R&D systems #S11150), 100 IU/mL penicillin, and 100 μg/mL streptomycin) (Life Technologies #15140). One day prior to infection, cells were seeded in antibiotic-free media at 2×10^4^ (for automated fluorescent microscopy assay) or 4×10^4^ cells/well (for real-time cytotoxicity assay) into 96-well tissue culture treated plates (Corning 3603 for plate reader, Greiner μClear #655096 for automated microscopy assays) and when indicated treated with 10 ng/ml IFNγ overnight (0.5% BSA, Peprotech) or disulfiram (30 μM) in DMSO (Cayman Chemical) one hour prior to and throughout the infection.

#### Knockout cell lines

All mammalian knockout cells were generated via CRISPR/Cas9 editing. To target GSDMD, a single sgRNA (sgRNA-GSDMD1)-Cas9 RNP complex was used (Synthego), while to target GBP1, a mixture of three sgRNAs (sgRNA-GBP1-1, sgRNA-GBP1-2, and sgRNA-GBP1-3) in complex with Cas9 was used (Gene Knockout Kit v2) (Synthego). Guide sequences are defined in Table S3. In each case, the RNP complexes were electroporated into 1×10^6^ cells (HeLa or HCT8) using a Neon system (Thermo Fisher Scientific). Pools were monitored for knockout efficiency after a 2–3-day recovery period. Cells were diluted to isolate single cells, which were expanded to generate clonal cell lines. Knockouts were verified by lysing 1×10^6^ cells seeded in wells of 6-well plates for 15 min with 250 μl ice-cold radioimmunoprecipitation assay (RIPA) buffer (25 mM Tris, pH 8, 150 mM NaCl, 0.1% SDS, 1% NP-40 plus Complete cocktail protease inhibitors (Millipore Sigma #11836170001). A cell scraper was used to remove lysed cells from the plate, which was transferred to a 1.5 ml tube and centrifuged at 13,000g for 15 min. The supernatant was resuspended in protein-loading dye, and samples were separated by SDS-PAGE, transferred to nitrocellulose membranes, and immunoblotted with antibodies using conditions described in Table S4.

#### Bacterial infections

For *Shigella* infections, single Congo red colonies were picked and grown overnight in 2 ml TCS (tryptic soy broth) at 30°C with aeration. For mT3Ec infections, single colonies were picked and grown overnight in 2 ml TCS at 37°C with aeration. The next day, each culture was back diluted 1:100 into TCS and grown at 37°C with aeration for 2-3 h until the cultures reached an OD_600_ of 0.8-1.0. For mT3Ec and in cases where IpaH9.8 expression was induced in *Shigella*, IPTG at a final concentration of 1 mM was added to the media at 1 h post-back dilution. After a total of 2-3 h post back dilution, when the cultures reached an OD_600_ of 0.8-1.0, bacteria were pelleted and resuspended in prewarmed low-glucose DMEM (no phenol red, Invitrogen #11054-020) plus 1% FBS then diluted to the indicated MOI for infections. In parallel, cell lines were washed 3 times with the same DMEM. Bacteria were added to each well at designated MOI, and the plates were centrifuged at 2000 RPM for 10 min to synchronize infection. The plates were incubated at 37°C in a 5% CO_2_ incubator for 30 min, after which each well was washed three times with HBSS/10% FBS/50 mM HEPES containing gentamicin (50 μg/mL) to kill and remove extracellular bacteria.

#### Plate reader-based cell death assay

Infections were carried out as described above in 96-well clear bottom tissue culture-treated plates (see above). During the washing step, propidium iodide (PI) was added to a final concentration of 3 μM to identify cells undergoing death via pyroptosis. After the washes, 200 μl of HBSS/FBS/HEPES/GENT was added to each well. PI uptake of infected cells was monitored using a SpectraMax i3x plate reader (Molecular Devices). A 3 × 3 grid of fluorescent output reads covering each well was taken and averaged every 10 min over a defined time course.

#### Automated fluorescent microscopy assay

Infections were carried out as described above in 96-well clear bottom tissue culture treated plates (see above). During the washing step, propidium iodide (PI) was added to a final concentration of 3 μM to identify cells undergoing death via pyroptosis, 16.2 μM Hoechst 33342 to identify all cells, and when indicated, 0.2% arabinose to monitor the growth of intracellular *Shigella* via sfGFP-expression. The plate was imaged using an ImageXpressPICO automated microscope (Molecular Devices) using a 20x/0.4 lens with DAPI, Texas Red, and, when indicated, FITC filters. Images were obtained every 10-15 min. Images of each well were analyzed using the Cell Reporter Xpress software (Molecular Devices). The percentage of dead cells was calculated by dividing the total number of PI-positive cells by the total number of Hoechst-stained cells imaged in each well. Similarly, the percentage of invaded cells was calculated by dividing the total number of GFP-positive cells by the total number of Hoechst-stained cells imaged in each well. GFP fluorescence was calculated by dividing the total GFP area by the number of Hoechst-stained cells imaged in each well.

In each case, the results from three replicate wells were averaged.

#### Secretion assay

Overnight cultures grown in TCS were diluted 1:100 into 2 ml of TCS and incubated at 37°C with aeration. One mM IPTG was added when cultures reached OD_600_ of ~0.4. After two hours, when mid-log growth was observed, equivalent numbers of bacteria from each culture were pelleted, resuspended in 2 ml of phosphate-buffered saline (PBS) supplemented with 10 μM Congo red (Sigma), and incubated for 30 min. Bacterial cultures were centrifuged, and the cell pellets were resuspended in protein-loading dye. Proteins in the supernatant fractions were TCA (trichloroacetic acid)-precipitated [10% (vol/vol)] and resuspended in loading dye. Proteins from supernatant fractions and cell lysates were separated via SDS-PAGE and immunoblotted with designated antibodies.

#### Cell lysates of infected cells

HeLa cells seeded in 6-well plates (1×10^6^cells/well) were infected as described above. Two hours post-infection, each well was washed three times with pre-warmed PBS, after which the cells were incubated for 15 min on ice in 250 μl RIPA buffer, which does not lyse bacterial cells. A cell scraper was used to remove lysed cells from the plate, and the mixture was centrifuged at 13,000g for 15 min. The supernatant was resuspended in protein-loading dye, and samples were separated by SDS-PAGE, transferred to nitrocellulose membranes, and immunoblotted with antibodies using conditions described in Table S4.

#### Intracytosolic LPS Assay

HeLa cells were seeded in 6-well plates at a density of 1×10^6^ cells/well and when indicated, treated with 10 ng/ml of IFNγ overnight. In the morning, cells were infected with *Shigella* (MOI of 3), as described above. At 2 hours post-infection, cells were washed three times with prewarmed PBS before being treated with 500 μL prewarmed TrypLE Trypsin (Life Technologies #12605). After a 5-10 min incubation at 37°C, the cells were resuspended in 500 μL prewarmed DMEM with 10% FBS and transferred to 1.5 ml Eppendorf tubes which were spun at 100g for 10 min. The supernatants were discarded, and pellets were resuspended in 500 μL ice-cold 0.005% digitonin in PBS and incubated at 4°C for 10 min on a rotator. The lysed cells were centrifuged at 400g for 10 min to separate soluble and residual fractions. The supernatant (cytosolic) fractions (200 μl) were applied to AcroPrep Advance 96-well Filter Plates (0.2 μm, polyethersulfone membrane, Pall Corp, #8019) to remove intact bacteria. Filtered cytosolic fractions were diluted 1:100 in endotoxin-free water, after which LPS was quantified using a Limulus amoebocyte lysate (LAL) chromogenic assay according to the manufacturer’s instructions (Associates of Cape Cod #E0005 and #C0031). The pellet (residual) fractions were each resuspended in 500 μL of 0.1% CHAPS in HBSS and incubated at RT for 10 min. Samples were immunoblotted with designated antibodies to confirm fractionation and assess GBP1 degradation. When comparing specific variables, we quantified all samples in parallel, as significant differences in absolute, but not relative, cLPS levels were detected on different days with the same samples, likely due to the sensitivity of the assay in detecting minor alterations in LPS standard levels. To quantify the relative numbers of infecting bacteria per condition, the residual fraction for each sample was serially diluted, plated, and colony-forming units (CFU) were quantified using ImageJ.

#### Statistical analyses

Prism 9.4.1 (GraphPad Software) was used to graph data and conduct all statistical analyses. Differences were considered statistically significant if the p-value was < 0.05.

## Acknowledgments

We thank Cerina Karr and Elizabeth Turcotte for assistance in generating knockout cell lines and Drs. Ann Hochschild, Thomas Bernhardt, and Marcia Goldberg for supplying plasmids. Drs. Russell Vance and Miriam Kutsch for critically reading the manuscript. This work was supported by National Institute of Health grants AI064285 (C.F.L), AI28360 (C.F.L.), AI007061 (K.K.), and AI139425 (JC).

## References

1. X. Liu, J. Lieberman, A Mechanistic Understanding of Pyroptosis: The Fiery Death Triggered by Invasive Infection. Adv Immunol 135, 81–117 (2017).

2. J. A. Hagar, D. A. Powell, Y. Aachoui, R. K. Ernst, E. A. Miao, Cytoplasmic LPS activates caspase-11: implications in TLR4-independent endotoxic shock. Science 341, 1250–1253 (2013).

3. N. Kayagaki et al., Noncanonical inflammasome activation by intracellular LPS independent of TLR4. Science 341, 1246–1249 (2013).

4. J. Shi et al., Inflammatory caspases are innate immune receptors for intracellular LPS. Nature 514, 187–192 (2014).

5. N. Kayagaki et al., Caspase-11 cleaves gasdermin D for non-canonical inflammasome signalling. Nature 526, 666–671 (2015).

6. J. Shi et al., Cleavage of GSDMD by inflammatory caspases determines pyroptotic cell death. Nature 526, 660–665 (2015).

7. X. Liu et al., Inflammasome-activated gasdermin D causes pyroptosis by forming membrane pores. Nature 535, 153–158 (2016).

8. X. Mou, S. Souter, J. Du, A. Z. Reeves, C. F. Lesser, Synthetic bottom-up approach reveals the complex interplay of Shigella effectors in regulation of epithelial cell death. Proc Natl Acad Sci U S A 115, 6452–6457 (2018).

9. N. Naseer et al., Salmonella enterica Serovar Typhimurium Induces NAIP/NLRC4- and NLRP3/ASC-Independent, Caspase-4-Dependent Inflammasome Activation in Human Intestinal Epithelial Cells. Infect Immun 90, e0066321 (2022).

10. M. P. Wandel et al., Guanylate-binding proteins convert cytosolic bacteria into caspase-4 signaling platforms. Nat Immunol 21, 880–891 (2020).

11. L. A. Knodler et al., Noncanonical inflammasome activation of caspase-4/caspase-11 mediates epithelial defenses against enteric bacterial pathogens. Cell Host Microbe 16, 249–256 (2014).

12. R. Finethy et al., Inflammasome Activation by Bacterial Outer Membrane Vesicles Requires Guanylate Binding Proteins. mBio 8(2017).

13. B. Lagrange et al., Human caspase-4 detects tetra-acylated LPS and cytosolic Francisella and functions differently from murine caspase-11. Nat Commun 9, 242 (2018).

14. D. M. Pilla et al., Guanylate binding proteins promote caspase-11-dependent pyroptosis in response to cytoplasmic LPS. Proc Natl Acad Sci U S A 111, 6046–6051 (2014).

15. J. C. Santos et al., LPS targets host guanylate-binding proteins to the bacterial outer membrane for non-canonical inflammasome activation. EMBO J 37(2018).

16. M. Kutsch et al., Direct binding of polymeric GBP1 to LPS disrupts bacterial cell envelope functions. EMBO J 39, e104926 (2020).

17. J. C. Santos et al., Human GBP1 binds LPS to initiate assembly of a caspase-4 activating platform on cytosolic bacteria. Nat Commun 11, 3276 (2020).

18. S. Zhu et al., Cryo-ET of a human GBP coatomer governing cell-autonomous innate immunity to infection. bioRxiv 10.1101/2021.08.26.457804, 2021.2008.2026.457804 (2021).

19. P. Li et al., Ubiquitination and degradation of GBPs by a Shigella effector to suppress host defence. Nature 551, 378–383 (2017).

20. A. S. Piro et al., Detection of Cytosolic Shigella flexneri via a C-Terminal Triple-Arginine Motif of GBP1 Inhibits Actin-Based Motility. mBio 8(2017).

21. M. P. Wandel et al., GBPs Inhibit Motility of Shigella flexneri but Are Targeted for Degradation by the Bacterial Ubiquitin Ligase IpaH9.8. Cell Host Microbe 22, 507–518 e505 (2017).

22. D. Fisch et al., Human GBP1 Differentially Targets Salmonella and Toxoplasma to License Recognition of Microbial Ligands and Caspase-Mediated Death. Cell Rep 32, 108008 (2020).

23. I. A. Khalil et al., Morbidity and mortality due to shigella and enterotoxigenic Escherichia coli diarrhoea: the Global Burden of Disease Study 1990-2016. Lancet Infect Dis 18, 1229–1240 (2018).

24. K. L. Kotloff, M. S. Riddle, J. A. Platts-Mills, P. Pavlinac, A. K. M. Zaidi, Shigellosis. Lancet 391, 801–812 (2018).

25. T. Kobayashi et al., The Shigella OspC3 effector inhibits caspase-4, antagonizes inflammatory cell death, and promotes epithelial infection. Cell Host Microbe 13, 570–583 (2013).

26. C. Oh, A. Verma, M. Hafeez, B. Hogland, Y. Aachoui, Shigella OspC3 suppresses murine cytosolic LPS sensing. iScience 24, 102910 (2021).

27. Z. Li et al., Shigella evades pyroptosis by arginine ADP-riboxanation of caspase-11. Nature 599, 290–295 (2021).

28. P. Clerc, P. J. Sansonetti, Entry of Shigella flexneri into HeLa cells: evidence for directed phagocytosis involving actin polymerization and myosin accumulation. Infect Immun 55, 2681–2688 (1987).

29. J. Du et al., The type III secretion system apparatus determines the intracellular niche of bacterial pathogens. Proc Natl Acad Sci U S A 113, 4794–4799 (2016).

30. M. Mavris, P. J. Sansonetti, C. Parsot, Identification of the cis-acting site involved in activation of promoters regulated by activity of the type III secretion apparatus in Shigella flexneri. J Bacteriol 184, 6751–6759 (2002).

31. H. Ashida et al., A bacterial E3 ubiquitin ligase IpaH9.8 targets NEMO/IKKgamma to dampen the host NF-kappaB-mediated inflammatory response. Nat Cell Biol 12, 66–73; sup pp 61-69 (2010).

32. J. Okuda et al., Shigella effector IpaH9.8 binds to a splicing factor U2AF(35) to modulate host immune responses. Biochem Biophys Res Commun 333, 531–539 (2005).

33. C. Ji et al., Structural mechanism for guanylate-binding proteins (GBPs) targeting by the Shigella E3 ligase IpaH9.8. PLoS Pathog 15, e1007876 (2019).

34. S. K. Vanaja et al., Bacterial Outer Membrane Vesicles Mediate Cytosolic Localization of LPS and Caspase-11 Activation. Cell 165, 1106–1119 (2016).

35. M. Ramsby, G. Makowski, Differential detergent fractionation of eukaryotic cells. Cold Spring Harb Protoc 2011, prot5592 (2011).

36. J. J. Hu et al., FDA-approved disulfiram inhibits pyroptosis by blocking gasdermin D pore formation. Nat Immunol 21, 736–745 (2020).

37. M. S. Dickinson et al., GBP2 aggregates LPS and activates the caspase-4 inflammasome independent of the bacterial encapsulation factor GBP1. bioRxiv 10.1101/2022.10.05.511023, 2022.2010.2005.511023 (2022).

