## Supplemental FIgures for "*Shigella* IpaH9.8 limits GBP1-dependent LPS release from intracytosolic bacteria to suppress caspase-4 activation"

### Figure S1

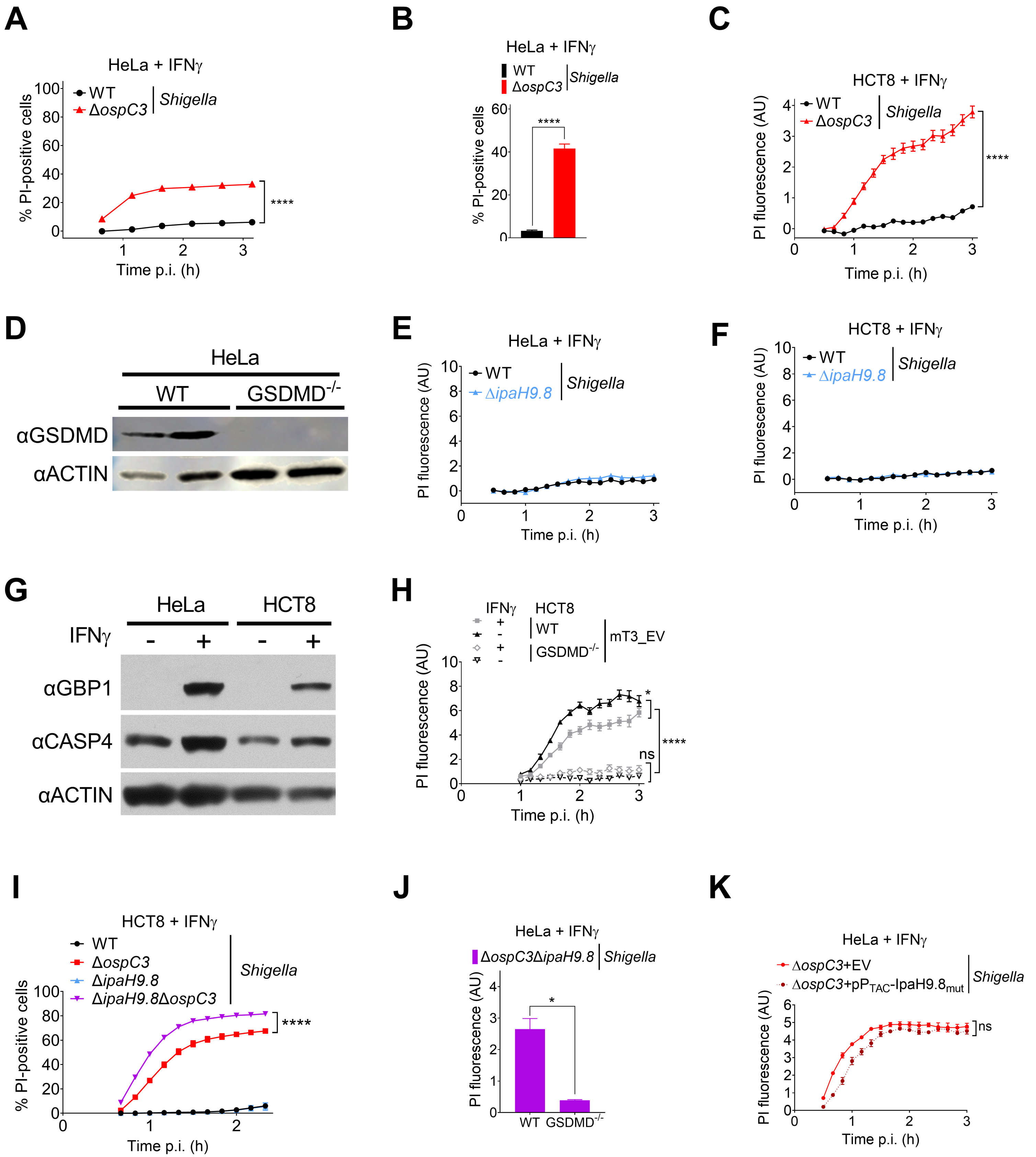
A
HeLa + IFNγ
WT
ΔospC3
Shigella
% PI-positive cells
Time p.i. (h)
****
B
HeLa + IFNγ
WT
ΔospC3
Shigella
% PI-positive cells
****
C
HCT8 + IFNγ
WT
ΔospC3
Shigella
PI fluorescence (AU)
Time p.i. (h)
****
D
HeLa
WT
GSDMD-/-
αGSDMD
αACTIN
E
HeLa + IFNγ
WT
ΔipaH9.8
Shigella
PI fluorescence (AU)
Time p.i. (h)
F
HCT8 + IFNγ
WT
ΔipaH9.8
Shigella
PI fluorescence (AU)
Time p.i. (h)
G
HeLa
HCT8
IFNγ
αGBP1
αCASP4
αACTIN
H
IFNγ
HCT8
WT
GSDMD-/-
mT3_EV
PI fluorescence (AU)
Time p.i. (h)
*
ns
****
I
HCT8 + IFNγ
WT
ΔospC3
ΔipaH9.8
ΔipaH9.8ΔospC3
Shigella
% PI-positive cells
Time p.i. (h)
****
J
HeLa + IFNγ
ΔospC3ΔipaH9.8
Shigella
PI fluorescence (AU)
WT
GSDMD-/-
*
K
HeLa + IFNγ
ΔospC3+EV
ΔospC3+pPTAC-IpaH9.8mut
Shigella
PI fluorescence (AU)
Time p.i. (h)
ns

**Figure S1: *Shigella* OspC3 and IpaH9.8 cooperate to suppress bacterial-triggered pyroptosis of IFNγ-primed epithelial cells.** (A-C, E, F, H-K) WT or $GSDMD^{-/-}$ HeLa or HCT8 cells primed overnight with 10 ng/ml IFNγ were infected with the indicated *Shigella* or mT3Ec strains at MOI of 100 (A) or 10 (B, C, E, F, H-K). Thirty minutes post-infection, infected cells were treated with gentamicin, PI, and in (A, B, I) with Hoechst. Cell death was assessed by monitoring $PI^+$/$Hoechst^+$ cells using an automated imaging system (A-B, I) or PI-uptake using a plate reader (C, E-F, H, J, K). Time courses of cell death (A, C, E-F, H-I, K) or 3-hr endpoint (B, J). Cells were infected with *Shigella* that carry pBAD33-AfaI in (C, E-F, I-K) and pNG162-AfaI in (B). Values shown are the mean +/- SEM of three experimental replicates. Where indicated, statistical significance was assessed using a two-way ANOVA with Tukey's post hoc test (A, C, I), a t-test (J), or an ordinary one-way ANOVA (B, H, K). *P < 0.05, **P < 0.01, ***P < 0.001, ****P < 0.0001, ns = non-significant. (D, G) Immunoblots of lysates of unprimed (D, G) and IFNγ-primed (G) HeLa (D, G) and HCT8 (G) cells immunoblotted with designated antibodies. Blots are representative of at least three independent experiments.

### Figure S2

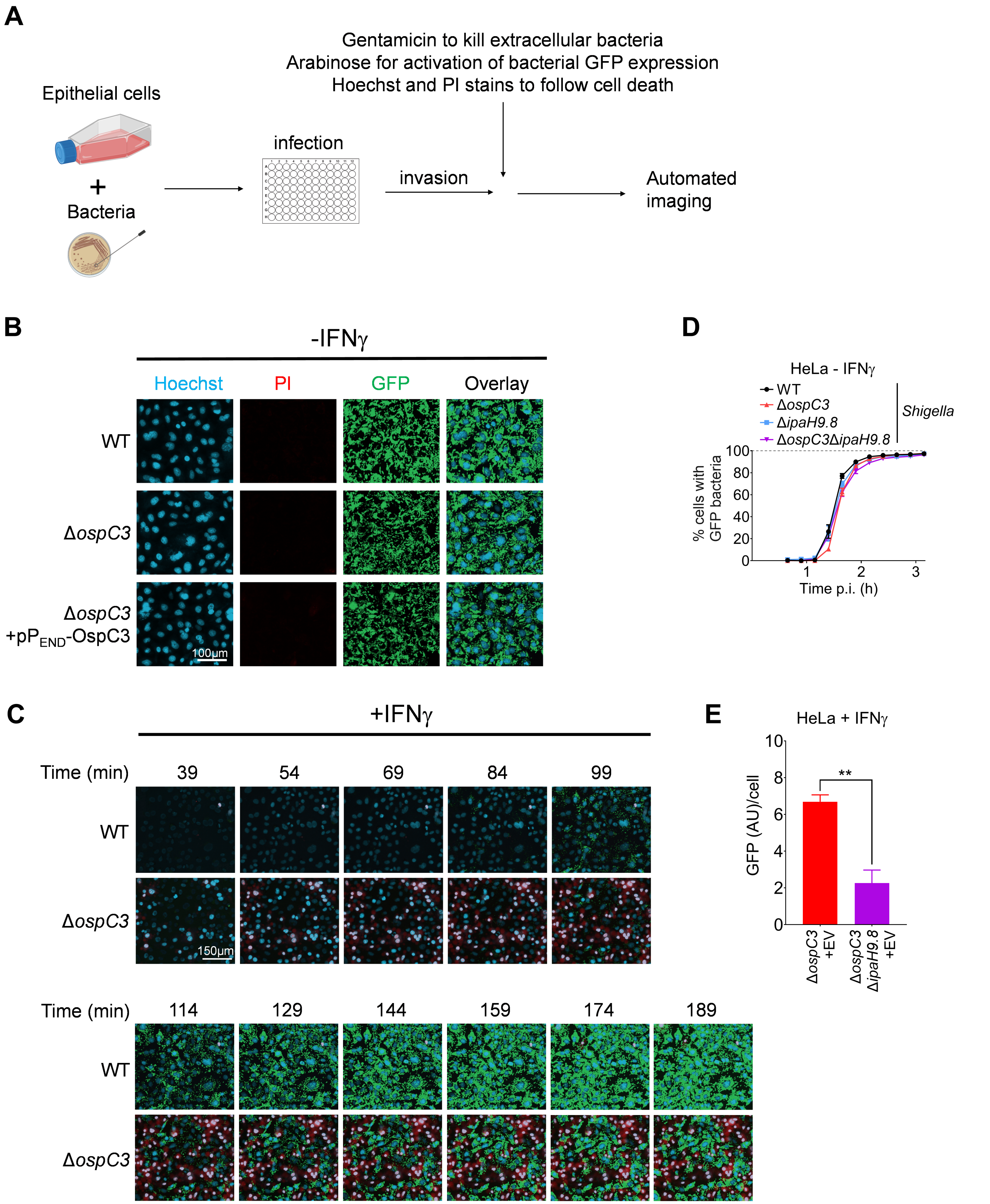

A
Epithelial cells
+
Bacteria
infection
invasion
Gentamicin to kill extracellular bacteria
Arabinose for activation of bacterial GFP expression
Hoechst and PI stains to follow cell death
Automated imaging
B
-IFNγ
Hoechst
PI
GFP
Overlay
WT
ΔospC3
ΔospC3 +pPEND-OspC3
100µm
C
+IFNγ
Time (min)
39
54
69
84
99
WT
ΔospC3
150µm
Time (min)
114
129
144
159
174
189
WT
ΔospC3
D
HeLa - IFNγ
WT
ΔospC3
ΔipaH9.8
ΔospC3ΔipaH9.8
Shigella
% cells with GFP bacteria
0
20
40
60
80
100
1
2
3
Time p.i. (h)
E
HeLa + IFNγ
GFP (AU)/cell
0
2
4
6
8
10
**
ΔospC3 +EV
ΔospC3 ΔipaH9.8 +EV

**Figure S2: *Shigella* OspC3 and IpaH9.8 cooperate to promote the growth of intracytosolic *Shigella*.** (A) Flow chart of the automated microscopy-based assay used to quantify cell death and replication of intracellular bacteria. Created with BioRender.com. (B-E) HeLa cells unprimed (B, D) or primed overnight with 10 ng/ml IFN$\gamma$ (C, E) were infected with designated strains that carry pBAD33-sfGFP, pBR322-AfaI and an empty vector (EV) at an MOI of 10. Thirty minutes post-infection, PI, Hoechst, and arabinose were added to the medium, and cells were imaged using an automated fluorescent microscope. (B-C) Representative images of cells stained with Hoechst (blue), PI (red), and GFP (green). (D) Time course of percentage cells containing GFP bacteria (grey line indicates 100%), and (E) 3-hour end point of GFP/Hoechst$^{+}$ cells. Values shown are the mean +/- SEM of three experimental replicates. Where indicated, statistical significance was assessed using a t-test (E). *$P < 0.05$, **$P < 0.01$, ***$P < 0.001$, ****$P < 0.0001$, ns = non-significant.

### Figure S3

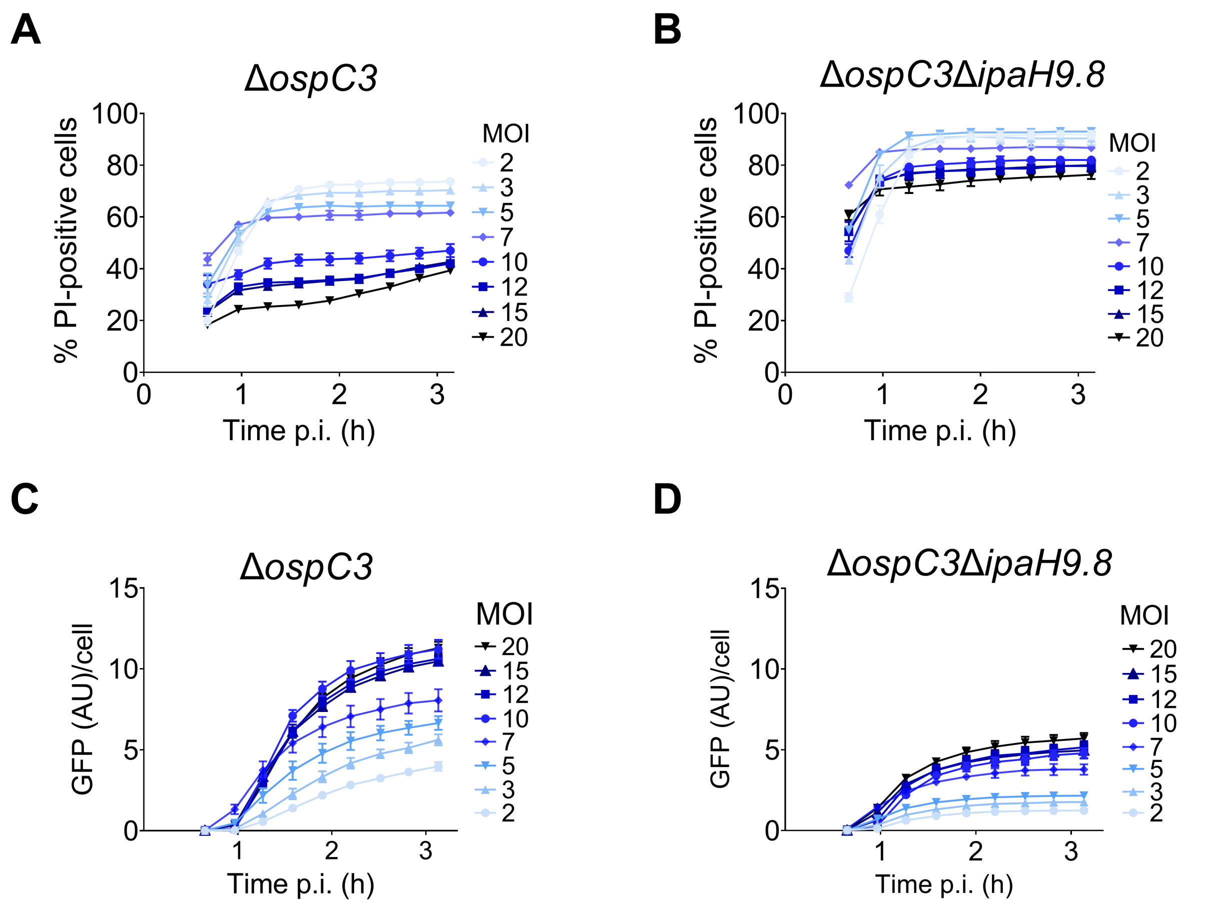
A
ΔospC3
% PI-positive cells
Time p.i. (h)
MOI
2
3
5
7
10
12
15
20
B
ΔospC3ΔipaH9.8
% PI-positive cells
Time p.i. (h)
C
ΔospC3
GFP (AU)/cell
Time p.i. (h)
D
ΔospC3ΔipaH9.8
GFP (AU)/cell
Time p.i. (h)

**Figure S3: Time course of PI-uptake and bacterial replication of cells infected with Δ*ospC3* or Δ*ospC3*Δ*ipaH9.8 Shigella.*** (A-B) WT HeLa cells primed overnight with 10 ng/ml IFNγ were infected with Δ*ospC3* or Δ*ospC3*Δ*ipaH9.8 Shigella* at MOIs ranging from 2-20. Thirty minutes post-infection, cells were treated with PI, Hoechst, and arabinose and then imaged using an automated fluorescent microscope. Time course of $PI^+$/$Hoechst^+$ cells infected with Δ*ospC3* (A) or Δ*ospC3*Δ*ipaH9.8 Shigella* (B). Time course of GFP/$Hoechst^+$ cells infected with Δ*ospC3* (C) or Δ*ospC3*Δ*ipaH9.8 Shigella* (D).

### Figure S4

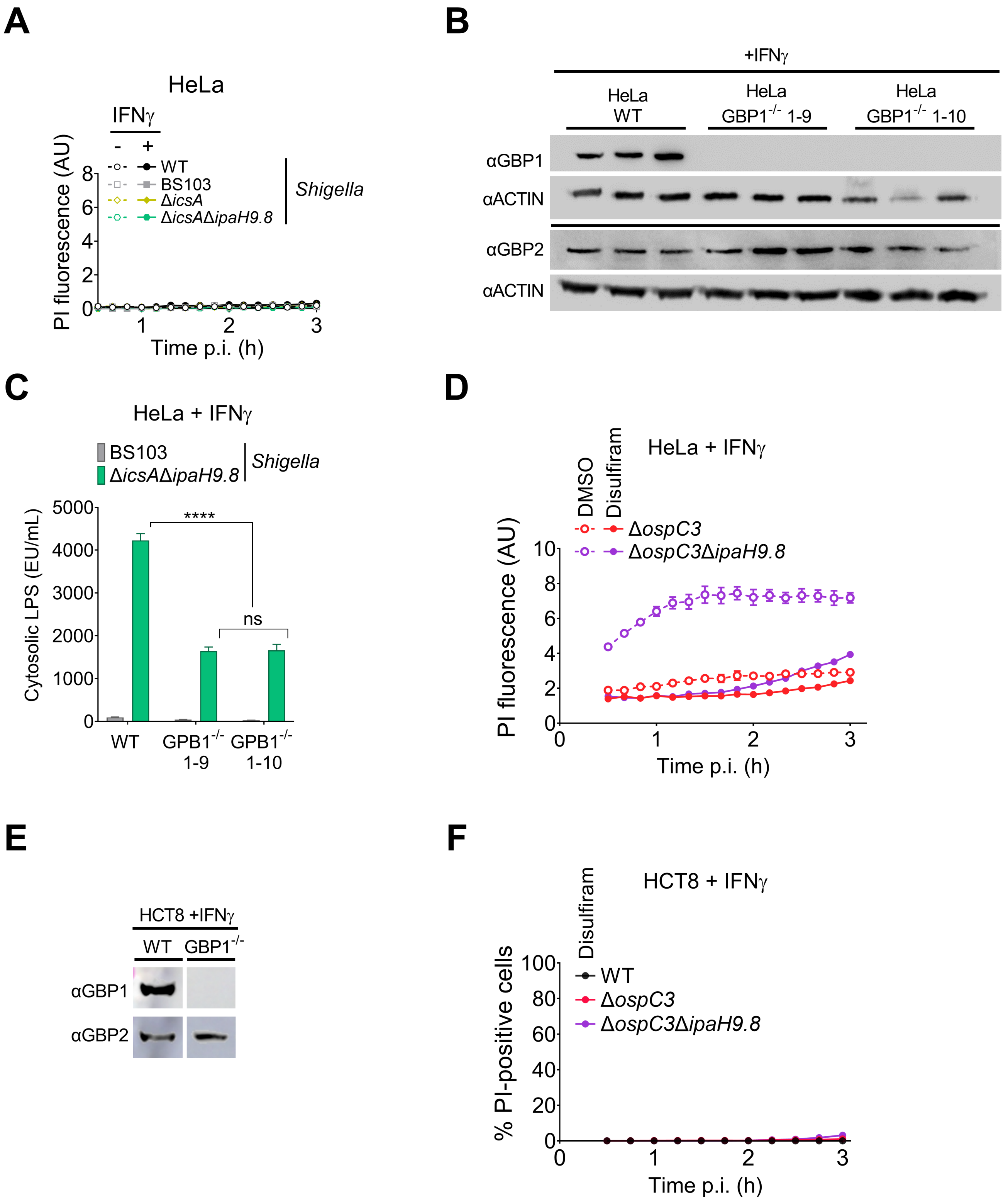

A
HeLa
IFNγ
- +
WT
BS103
ΔicsA
ΔicsAΔipaH9.8
Shigella
PI fluorescence (AU)
Time p.i. (h)
B
+IFNγ
HeLa WT
HeLa GBP1-/- 1-9
HeLa GBP1-/- 1-10
αGBP1
αACTIN
αGBP2
αACTIN
C
HeLa + IFNγ
BS103
ΔicsAΔipaH9.8
Shigella
Cytosolic LPS (EU/mL)
****
ns
WT
GPB1-/- 1-9
GPB1-/- 1-10
D
HeLa + IFNγ
DMSO
Disulfiram
ΔospC3
ΔospC3ΔipaH9.8
PI fluorescence (AU)
Time p.i. (h)
E
HCT8 +IFNγ
WT
GBP1-/-
αGBP1
αGBP2
F
HCT8 + IFNγ
Disulfiram
WT
ΔospC3
ΔospC3ΔipaH9.8
% PI-positive cells
Time p.i. (h)

**Figure S4: GBP1 promotes LPS release from intracellular *Shigella* and is non-essential for *Shigella*-triggered pyroptosis.** (A, C, D, F) WT HeLa cells unprimed and primed with 10 ng/ml IFNγ overnight were infected with designated strains that carry pNG162-AfaI at an MOI of 3. Thirty minutes p.i., PI was added to the medium, and cell death was monitored by PI-uptake using a plate reader (A, D), or PI and Hoechst were added, and cell death was assessed by monitoring $PI^+$/$Hoechst^+$ cells using an automated imaging system (F). When indicated, cells were pretreated with DMSO or DMSO/disulfiram, which was maintained in the medium throughout the infection. (B, E) Lysates of IFNγ-primed WT and $GBP1^{-/-}$ HeLa and HCT8 cells immunoblotted with designated antibodies. Images shown in E are each cropped from a single image (C) Quantification of cLPS levels in lysates of WT and $GBP1^{-/-}$ HeLa cells infected with BS103 or Δ*icsA*Δ*ipaH9.8 Shigella* that carry pNG162-AfaI at an MOI of 3. Values shown are the mean +/- SEM of three experimental replicates. Three biological replicates were performed, and representative data are shown. Data were analyzed using two-way ANOVA with Tukey's post hoc test. *P < 0.05, **P < 0.01, ***P < 0.001, ****P < 0.0001, ns = non-significant.

**Figure S1: *Shigella* OspC3 and IpaH9.8 cooperate to suppress bacterial-triggered pyroptosis of IFNγ-primed epithelial cells.** (A-C, E, F, H-K) WT or GSDMD$^{-/-}$ HeLa or HCT8 cells primed overnight with 10 ng/ml IFNγ were infected with the indicated *Shigella* or mT3Ec strains at MOI of 100 (A) or 10 (B, C, E, F, H-K). Thirty minutes post-infection, infected cells were treated with gentamicin, PI, and in (A, B, I) with Hoechst. Cell death was assessed by monitoring PI$^{+}$/Hoechst$^{+}$ cells using an automated imaging system (A-B, I) or PI-uptake using a plate reader (C, E-F, H, J, K). Time courses of cell death (A, C, E-F, H-I, K) or 3-hr endpoint (B, J). Cells were infected with *Shigella* that carry pBAD33-AfaI in (C, E-F, I-K) and pNG162-AfaI in (B). Values shown are the mean +/- SEM of three experimental replicates. Where indicated, statistical significance was assessed using a two-way ANOVA with Tukey's post hoc test (A, C, I), a t-test (J), or an ordinary one-way ANOVA (B, H, K). *P < 0.05, **P < 0.01, ***P < 0.001, ****P < 0.0001, ns = non-significant. (D, G) Immunoblots of lysates of unprimed (D, G) and IFNγ-primed (G) HeLa (D, G) and HCT8 (G) cells immunoblotted with designated antibodies. Blots are representative of at least three independent experiments.

### Figure S2

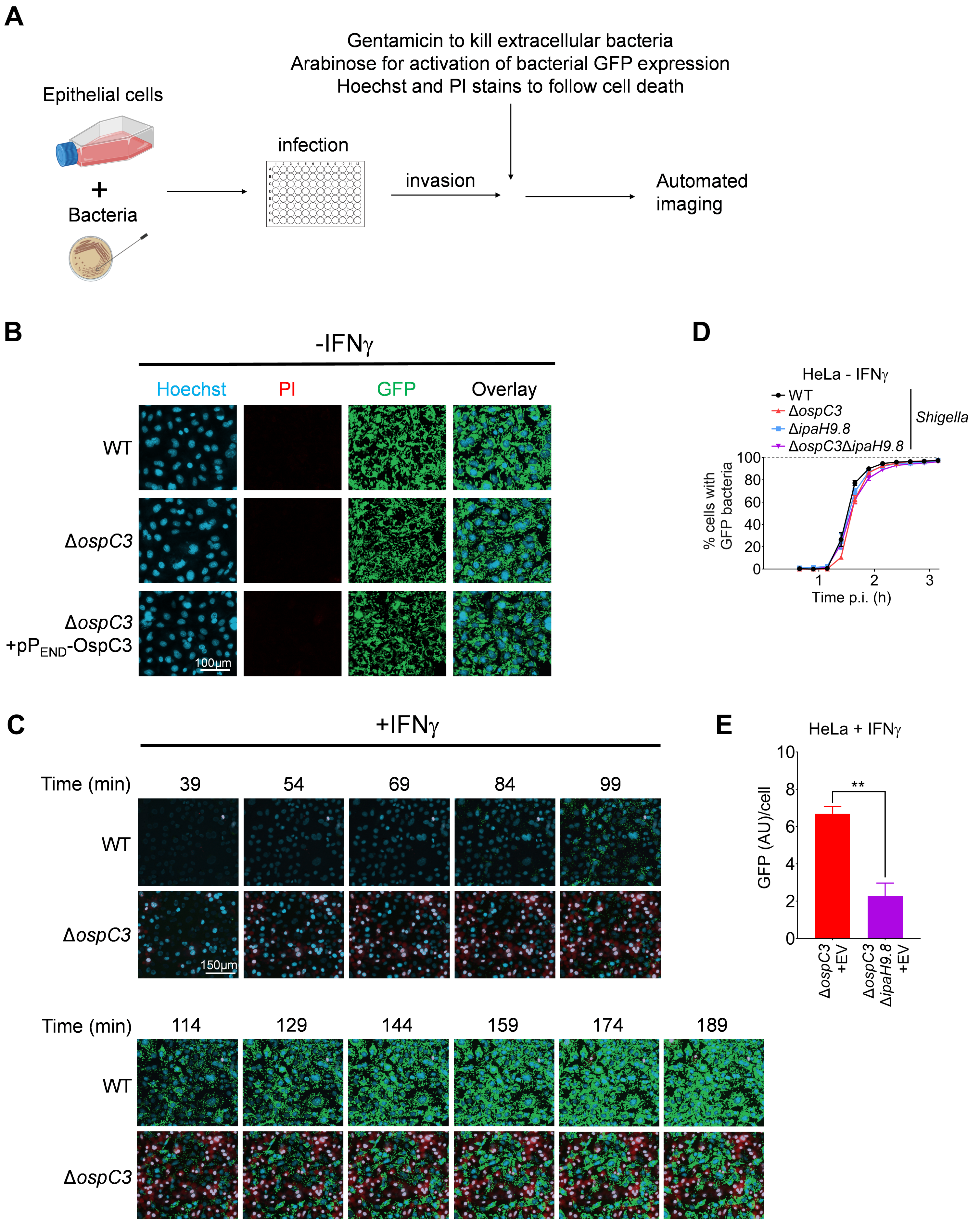

A
Epithelial cells
+
Bacteria
infection
invasion
Gentamicin to kill extracellular bacteria
Arabinose for activation of bacterial GFP expression
Hoechst and PI stains to follow cell death
Automated imaging
B
-IFNγ
Hoechst
PI
GFP
Overlay
WT
ΔospC3
ΔospC3 +pPEND-OspC3
100µm
C
+IFNγ
Time (min)
39
54
69
84
99
WT
ΔospC3
150µm
Time (min)
114
129
144
159
174
189
WT
ΔospC3
D
HeLa - IFNγ
WT
ΔospC3
ΔipaH9.8
ΔospC3ΔipaH9.8
Shigella
% cells with GFP bacteria
Time p.i. (h)
E
HeLa + IFNγ
GFP (AU)/cell
**
ΔospC3 +EV
ΔospC3 ΔipaH9.8 +EV

**Figure S2: *Shigella* OspC3 and IpaH9.8 cooperate to promote the growth of intracytosolic *Shigella*.** (A) Flow chart of the automated microscopy-based assay used to quantify cell death and replication of intracellular bacteria. Created with BioRender.com. (B-E) HeLa cells unprimed (B, D) or primed overnight with 10 ng/ml IFN$\gamma$ (C, E) were infected with designated strains that carry pBAD33-sfGFP, pBR322-AfaI and an empty vector (EV) at an MOI of 10. Thirty minutes post-infection, PI, Hoechst, and arabinose were added to the medium, and cells were imaged using an automated fluorescent microscope. (B-C) Representative images of cells stained with Hoechst (blue), PI (red), and GFP (green). (D) Time course of percentage cells containing GFP bacteria (grey line indicates 100%), and (E) 3-hour end point of GFP/Hoechst$^{+}$ cells. Values shown are the mean +/- SEM of three experimental replicates. Where indicated, statistical significance was assessed using a t-test (E). *P < 0.05, **P < 0.01, ***P < 0.001, ****P < 0.0001, ns = non-significant.

### Figure S3

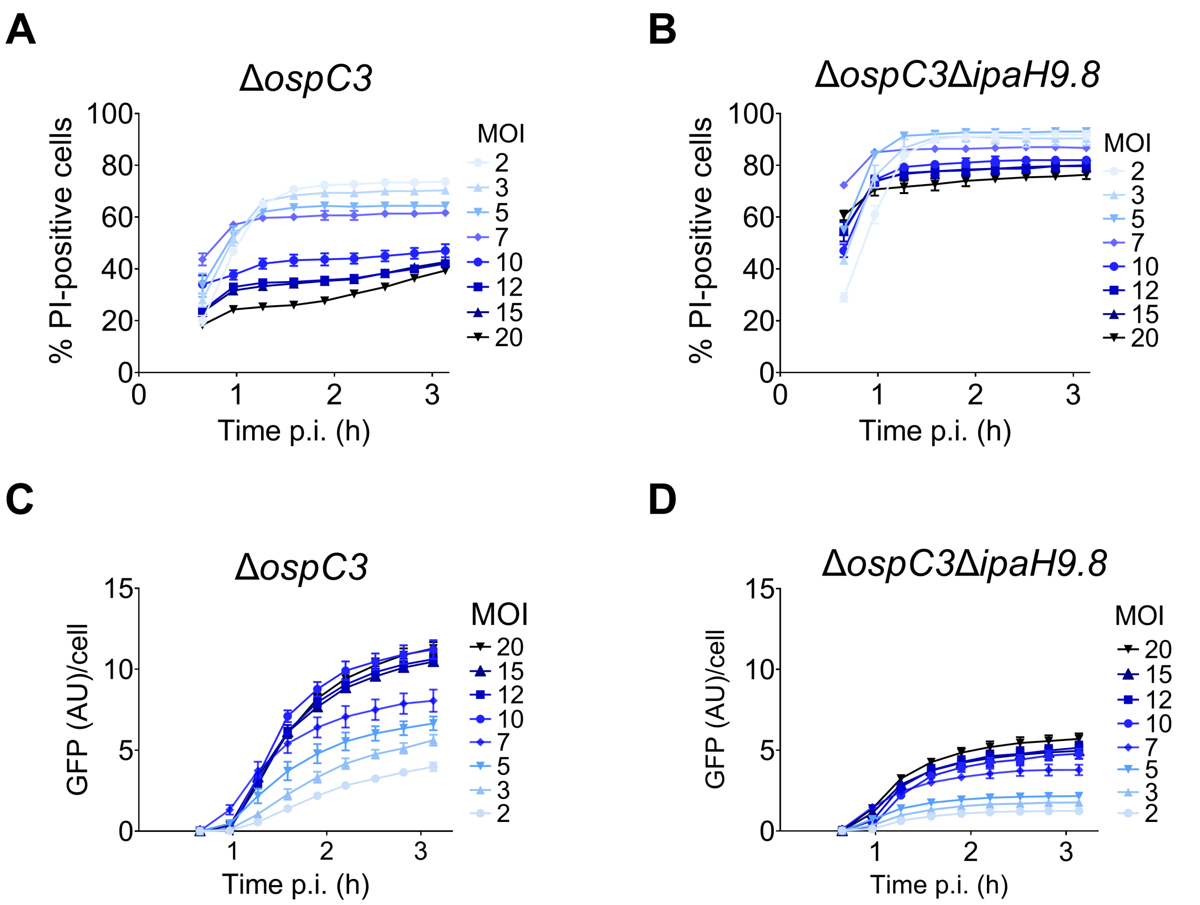
A
ΔospC3
% PI-positive cells
Time p.i. (h)
MOI
2
3
5
7
10
12
15
20
B
ΔospC3ΔipaH9.8
% PI-positive cells
Time p.i. (h)
MOI
C
ΔospC3
GFP (AU)/cell
Time p.i. (h)
MOI
D
ΔospC3ΔipaH9.8
GFP (AU)/cell
Time p.i. (h)
MOI

**Figure S3: Time course of PI-uptake and bacterial replication of cells infected with Δ*ospC3* or Δ*ospC3*Δ*ipaH9.8 Shigella.*** (A-B) WT HeLa cells primed overnight with 10 ng/ml IFNγ were infected with Δ*ospC3* or Δ*ospC3*Δ*ipaH9.8 Shigella* at MOIs ranging from 2-20. Thirty minutes post-infection, cells were treated with PI, Hoechst, and arabinose and then imaged using an automated fluorescent microscope. Time course of $PI^+$/$Hoechst^+$ cells infected with Δ*ospC3* (A) or Δ*ospC3*Δ*ipaH9.8 Shigella* (B). Time course of GFP/$Hoechst^+$ cells infected with Δ*ospC3* (C) or Δ*ospC3*Δ*ipaH9.8 Shigella* (D).

### Figure S4

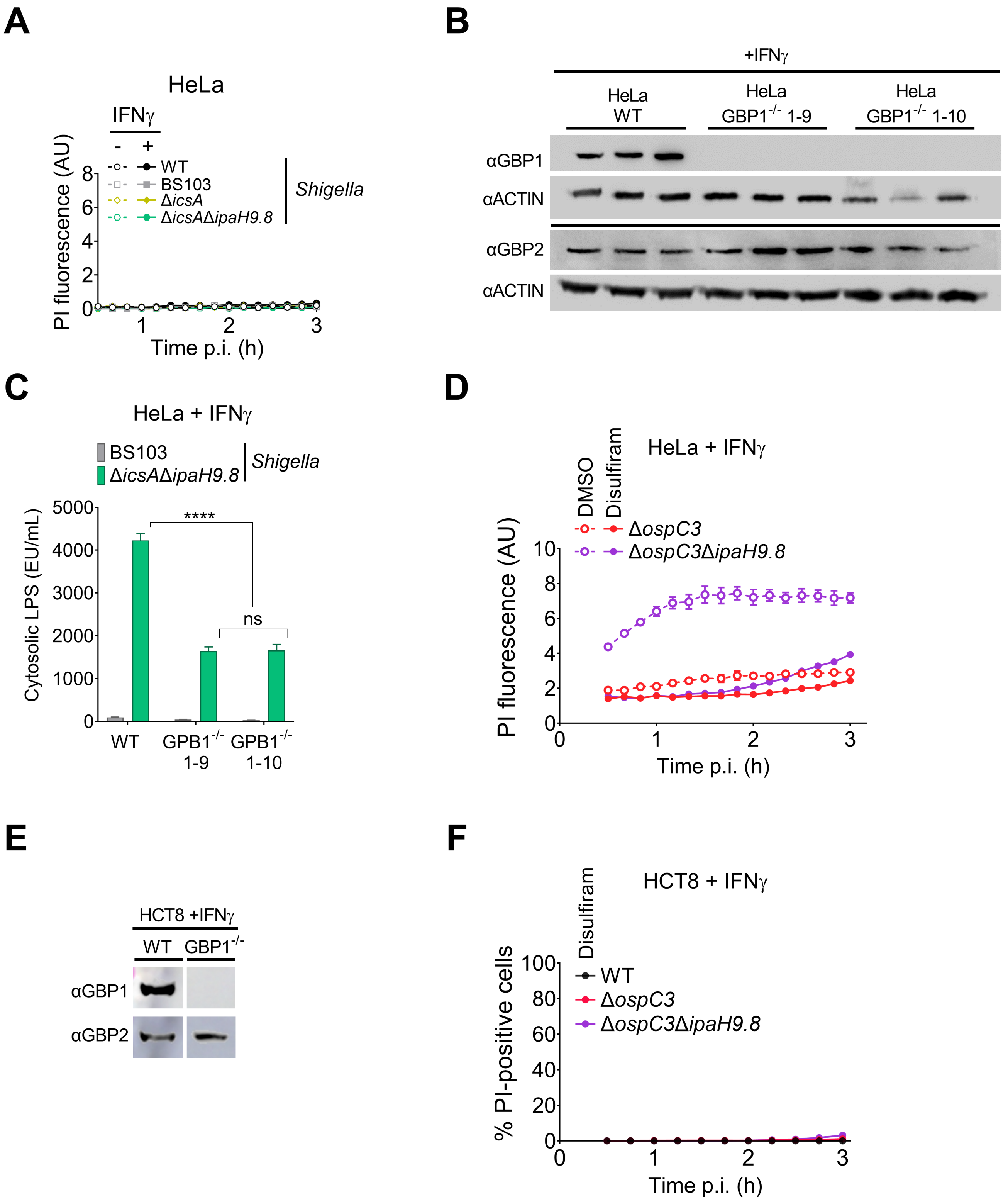
A
HeLa
IFNγ
- +
WT
BS103
ΔicsA
ΔicsAΔipaH9.8
Shigella
PI fluorescence (AU)
Time p.i. (h)
B
+IFNγ
HeLa WT
HeLa GBP1-/- 1-9
HeLa GBP1-/- 1-10
αGBP1
αACTIN
αGBP2
αACTIN
C
HeLa + IFNγ
BS103
ΔicsAΔipaH9.8
Shigella
Cytosolic LPS (EU/mL)
****
ns
WT
GPB1-/- 1-9
GPB1-/- 1-10
D
HeLa + IFNγ
DMSO
Disulfiram
ΔospC3
ΔospC3ΔipaH9.8
PI fluorescence (AU)
Time p.i. (h)
E
HCT8 +IFNγ
WT
GBP1-/-
αGBP1
αGBP2
F
HCT8 + IFNγ
Disulfiram
WT
ΔospC3
ΔospC3ΔipaH9.8
% PI-positive cells
Time p.i. (h)

**Figure S4: GBP1 promotes LPS release from intracellular *Shigella* and is non-essential for *Shigella*-triggered pyroptosis.** (A, C, D, F) WT HeLa cells unprimed and primed with 10 ng/ml IFNγ overnight were infected with designated strains that carry pNG162-AfaI at an MOI of 3. Thirty minutes p.i., PI was added to the medium, and cell death was monitored by PI-uptake using a plate reader (A, D), or PI and Hoechst were added, and cell death was assessed by monitoring $PI^+$/$Hoechst^+$ cells using an automated imaging system (F). When indicated, cells were pretreated with DMSO or DMSO/disulfiram, which was maintained in the medium throughout the infection. (B, E) Lysates of IFNγ-primed WT and $GBP1^{-/-}$ HeLa and HCT8 cells immunoblotted with designated antibodies. Images shown in E are each cropped from a single image (C) Quantification of cLPS levels in lysates of WT and $GBP1^{-/-}$ HeLa cells infected with BS103 or Δ*icsA*Δ*ipaH9.8 Shigella* that carry pNG162-AfaI at an MOI of 3. Values shown are the mean +/- SEM of three experimental replicates. Three biological replicates were performed, and representative data are shown. Data were analyzed using two-way ANOVA with Tukey's post hoc test. *P < 0.05, **P < 0.01, ***P < 0.001, ****P < 0.0001, ns = non-significant.
